# 3D analysis of the synaptic organization in the Entorhinal cortex in Alzheimer’s disease

**DOI:** 10.1101/2020.10.19.345025

**Authors:** M Domínguez-Álvaro, M Montero-Crespo, L Blazquez-Llorca, S Plaza-Alonso, N Cano-Astorga, J DeFelipe, L Alonso-Nanclares

## Abstract

The entorhinal cortex (EC) is especially vulnerable in the early stages of Alzheimer’s disease (AD). In particular, cognitive deficits have been linked to alterations in the upper layers of EC. In the present report, we examined layers II and III from eight human brain autopsies (four subjects with no recorded neurological alterations and four AD cases). We used stereological methods to assess cortical atrophy of the EC, and possible changes in the volume occupied by different cortical elements (neuronal and glial cell bodies; blood vessels; and neuropil). We performed 3D ultrastructural analyses of synapses using Focused Ion Beam/Scanning Electron Microscopy (FIB/SEM) to examine possible alterations related to AD.

At the light microscope level, we found a significantly lower volume fraction occupied by neuronal bodies in layer III and a higher volume fraction occupied by glial cell bodies in layer II in AD cases. At the ultrastructural level we observed that (i) there was a significantly lower synaptic density in both layers in AD cases; (ii) synapses were larger and more complex in layer II in AD cases; and (iii) there was a greater proportion of small and simple synapses in layer III in AD cases than in control individuals. These structural differences may play a role in the anatomical basis for the impairment of cognitive functions in AD.

**Significant Statement:** Analysis of the synaptic characteristics provides critical data on synaptic organization. Using 3D electron microscopy, the present study shows the synaptic organization of the neuropil of the human entorhinal cortex (EC) at the ultrastructural level. The EC is especially vulnerable in the early stages of Alzheimer’s disease (AD). Our present results show structural differences that may contribute as anatomical basis for the impairment of cognitive functions in AD. Thus, these results may help to understand the relationship between alterations of the synaptic circuits and the cognitive deterioration in AD.

## Introduction

Alzheimer’s disease (AD) is a progressive neurodegenerative disease which is considered to be the main cause of dementia. The brain of AD cases shows brain atrophy and, at the neuropathological level, the most characteristic findings include the presence of extracellular amyloid-β (Aβ) plaques and intracellular neurofibrillary tangles of filamentous aggregates of hyperphosphorylated tau protein (Alzheimer’s Association, 2020). Aβ plaques and neurofibrillary tangles are mostly found in the cerebral cortex, where both their numbers and the proportion of the cortex affected by them increase progressively as the disease advances (Braak and Braak, 1991; Dickson, 1997; Thal et al., 2002). Studies focusing on AD progression have shown that entorhinal cortex (EC) is one of the first brain regions affected by the presence of altered tau protein (Braak and Braak, 1991).

The EC has been shown to be essential for memory functions and spatial navigation (reviewed in Schultz et al., 2015), and alterations in its upper layers have been related to cognitive deficits in AD patients (Van Hoesen et al., 1991; Gomez-Isla et al., 1996). EC is considered to be an interface between the hippocampal formation and a large variety of association and limbic cortices (Lavenex and Amaral, 2000; Solodkin and Van Hoesen, 1996). In particular, the EC is the origin of the perforant pathway (from layers II and III), which provides the largest input source to the hippocampal formation, targeting the ammonic fields (CA) CA1, CA2 and CA3, as well as the dentate gyrus (DG) and subiculum. Specifically, layer II neurons project to CA1 via the trisynaptic circuit, passing through the dentate gyrus (DG) and CA3, while layer III neurons project directly to CA1 in the monosynaptic pathway (Insausti and Amaral, 2012; Kondo et al., 2009). It has been proposed that the trisynaptic pathway is more susceptible to premature degeneration (reviewed in Van Hoesen et al., 2006; Llorens-Martín et al., 2014).

The presence of pathological forms of Aβ and tau proteins in the cerebral cortex of AD cases has been related to neuronal loss, synapse alterations and dendritic spine degeneration (reviewed in Forner et al., 2017; Chen et al., 2019). The loss and dysfunction of synapses have been proposed as the major structural correlates of the cognitive decline associated with AD (Coleman et al., 2004; Dickson et al., 1995; Selkoe, 2002; Sze et al., 1997). It has been proposed that synaptic loss affects the subcortical regions and the entorhinal cortex first, and then progresses to other cortical regions (Braak and Braak, 1991; Braak and Del Tredici, 2012, 2020). Therefore, deciphering the changes that affect the normal function of synapses may contribute to better understanding of the pathological mechanisms of AD.

In the present study, we performed 3D ultrastructural analysis of the EC using Focused Ion Beam/Scanning Electron Microscopy (FIB/SEM) and specific software that allows the segmentation of synapses in a reconstructed 3D volume (Morales et al., 2011). This technology greatly facilitates the analysis of possible alterations at the synaptic level in AD, as previously shown in the human transentorhinal cortex and the CA1 hippocampal region (Domínguez-Álvaro et al., 2019; Montero-Crespo et al., 2021). Our goal was to investigate the possible synaptic changes occurring in layers II and III of the EC related to AD, not only with regard to numbers, spatial distribution and types of synapses, but also regarding the morphological characteristics of each synapse (shape and size), as well as possible changes in their postsynaptic targets. For this purpose, we performed an analysis of the neuropil from layers II and III of the EC, from eight human brain autopsies (four subjects with no recorded neurological alterations and four AD cases) with short postmortem delays (of less than 3.5h). Since the presence of Aβ plaques is related to a virtual lack of synapses in their vicinity (Blazquez-Llorca et al., 2013), we focused on the neuropil that was free of plaques. We also used stereological methods to assess cortical atrophy of the EC, and possible changes in the volume occupied by different cortical elements (neuronal and glial cell bodies, blood vessels and neuropil) at the light microscope level.

## Methods

### Tissue preparation

Human brain tissue was obtained from eight autopsies (4 male and 4 female subjects) with short postmortem delays (of less than 3.5 hours; supplied by Instituto de Neuropatología del IDIBELL-Hospital Universitario de Bellvitge, Barcelona, Spain; Unidad Asociada Neuromax, Laboratorio de Neuroanatomía Humana, Facultad de Medicina, Universidad de Castilla-La Mancha, Albacete and the Laboratorio Cajal de Circuitos Corticales UPM-CSIC, Madrid, Spain). The sampling procedure was approved by the Institutional Ethical Committees of each of the institutions involved. Tissue from some of these human brains has been used in previous studies (Domínguez-Álvaro et al., 2018, 2019, 2021; Montero-Crespo et al., 2020, 2021).

Briefly, tissue samples were obtained from 4 control cases (non-demented subjects with no recorded neurological or psychiatric alterations) and 4 AD cases according to the neuropathological criteria provided by the above-mentioned centers (Table 1).

**Table 1.** Clinical and neuropathological information. Braak Stages (Braak and Braak, 1991): III (NFTs —neurofibrillary tangles— in entorhinal cortex and closely related areas); III−IV (NFTs abundant in amygdala and hippocampus. Extending slightly into association cortex); V−VI (NFTs widely distributed throughout the neocortex and ultimately involving primary motor and sensory areas). CERAD Stages (Mirra et al., 1991): A (Low density of neuritic plaques); B (Intermediate density of neuritic plaques); C (High density of neuritic plaques). -: Not available.

| **Patient** | **Sex** | **Age**<br>**(years)** | **Cause of death** | **Postmortem**<br>**delay (h)** | **Braak**<br>**Stage** | **CERAD**<br>**Stage** | **Neuropsychological diagnosis** |
| --- | --- | --- | --- | --- | --- | --- | --- |
| **AB1** | Male | 45 | Lung cancer | <1 | 0 | 0 | No recorded neurological alterations |
| **AB3** | Male | 53 | Bladder carcinoma | 3.5 | 0 | 0 | No recorded neurological alterations |
| **IF10** | Male | 66 | Bronchopneumonia and cardiac failure | 2 | 0 | 0 | No recorded neurological alterations |
| **M16** | Male | 40 | Traffic accident | 3 | 0 | 0 | No recorded neurological alterations |
| **IF1** | Female | 80 | - | 2 | IV | B | No evidence of cognitive impairment and dementia |
| **VK11** | Female | 87 | Respiratory inflammation | 1.5 | III–IV | A | Dementia |
| **VK16** | Female | 88 | - | 2.5 | VI | C | Dementia |
| **VK22** | Female | 86 | - | 2 | V | C | Dementia |

Upon removal, the brain tissue was fixed in cold 4% paraformaldehyde (Sigma-Aldrich, St Louis, MO, USA) in 0.1M sodium phosphate buffer (PB; Panreac, 131965, Spain), pH 7.4, for 24-48h. After fixation, the tissue was washed in PB and coronally sectioned (150μm thick) in a vibratome (Vibratome Sectioning System, VT1200S Vibratome, Leica Biosystems, Germany). Sections containing EC were selected for Nissl staining, immunohistochemistry and electron microscopy (EM) processing (Fig. 1).

**Figure 1.**
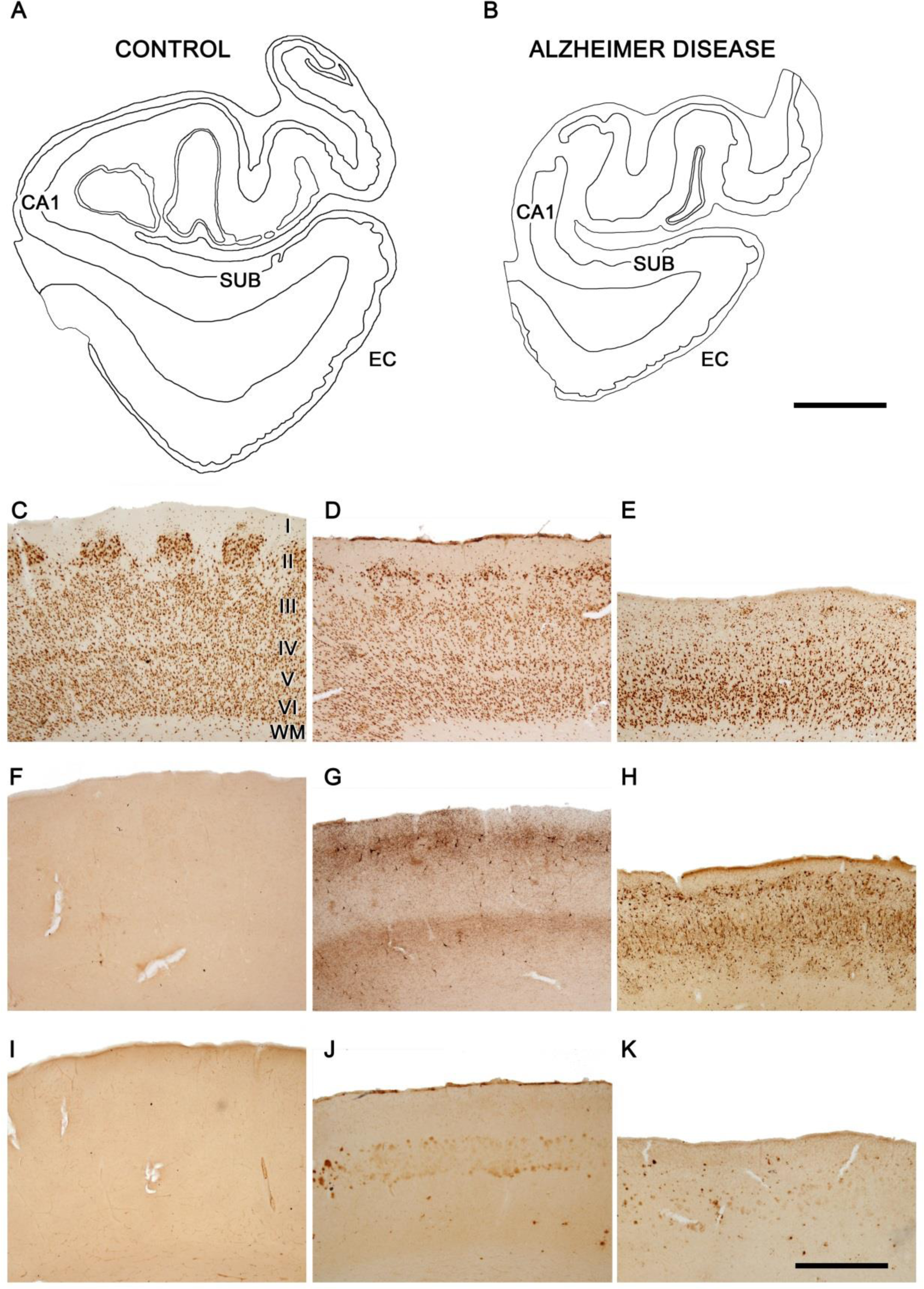
Entorhinal cortex of control and AD cases. Plots of the medial structures including the hippocampal formation and EC from control (**A**) and AD (**B**) cases to illustrate the atrophy that occurs in AD. Plots are based on Nissl-stained coronal sections from cases AB3 and IF1. Microphotographs showing the EC in a control case AB3 **(C, F, I)** and in AD cases IF1 **(D, G, J)** and VK16 **(E, H, K)**. Layers are indicated to the right of panel **C**. Sections are immunostained with antibodies anti-NeuN **(C-E)**, anti-PHF-Tau-AT8 **(F-H)** and anti-Aβ **(I-K)**. Note the intense labeling of PHF-Tau-AT8 positive neurons (G, H) and Aβ positive plaques (J, K) in AD cases, particularly in case VK16, and the lack of labeling in the control case. CA1: cornu ammonis field 1; EC: entorhinal cortex; SUB: subiculum; WM: white matter. Scale bar (in I): 4.4 mm in A, B and 1 mm in C−K.

Coronal sections of the human EC at medial level were used for the present study (reviewed in Insausti et al., 2017). The delimitation of the EC was established by combining the Nissl and anti-NeuN markers (Fig. 1). The main cytoarchitectural characteristic of the EC that allows it to be recognized is the presence of large islands of modified pyramidal neurons and stellate cells in layer II (Braak and Braak, 1992; Insausti and Amaral, 2012; Kobro-Flatmoen and Witter, 2019). Sections containing EC were selected for Nissl staining, immunohistochemistry and EM processing (Fig. 1).

### Immunohistochemistry

The selected sections were first rinsed in 0.1M PB, pretreated in 2% H_2_O_2_ for 30 minutes to remove endogenous peroxidase activity, and then incubated for 1h at room temperature in a solution of 3% normal horse serum (for monoclonal antibodies; Vector Laboratories Inc., Burlingame, CA) and 0.25% Triton-X (Merck, Darmstadt, Germany). The sections were then incubated for 48h at 4°C in the same solution with mouse anti-NeuN (1:2000; Chemicon; MAB377, Temecula, CA, USA) and anti-human PHF_-Tau_ antibody clone AT8 (1:2000, MN1020, Thermo Scientific, Waltham, MA, USA); for the sake of clarity, we will refer to this as anti-PHF_-Tau-AT8_. The sections selected for anti-Aβ were first treated with 88% formic acid (Sigma-Aldrich, No. 251364, St. Louis, MO, USA) to ensure specific plaque immunostaining, and were then incubated in a solution containing mouse antibody anti-Aβ (clone 6F/3D; 1:50, Dako M0872, Glostrup, Denmark). The sections were then processed with a secondary biotinylated horse anti-mouse IgG antibody (1:200, Vector Laboratories, Burlingame, CA, USA), and then incubated for 1h in an avidin-biotin peroxidase complex (Vectastain ABC Elite PK6100, Vector) and, finally, with the chromogen 3,3′-diaminobenzidine tetrahydrochloride (DAB; Sigma-Aldrich, St. Louis, MO, USA). Finally, the sections were dehydrated, cleared with xylene and cover-slipped.

### Tissue processing for EM

EC sections were postfixed for 24h in a solution containing 2% paraformaldehyde, 2.5% glutaraldehyde (TAAB, G002, UK) and 0.003% CaCl2 (Sigma, C-2661-500G, Germany) in sodium cacodylate (Sigma, C0250-500G, Germany) buffer (0.1M). These sections were washed in sodium cacodylate buffer (0.1M) and treated with 1% OsO4 (Sigma, O5500, Germany), 0.1% potassium ferrocyanide (Probus, 23345, Spain) and 0.003% CaCl2 in sodium cacodylate buffer (0.1M) for 1h at room temperature. After washing in PB, the sections were stained with 2% uranyl acetate (EMS, 8473, USA), and then dehydrated and flat-embedded in Araldite (TAAB, E021, UK) for 48h at 60°C (DeFelipe and Fairén, 1993). Embedded sections were glued onto a blank Araldite block and trimmed. Semithin sections (1–2 μm thick) were obtained from the surface of the block and stained with 1% toluidine blue (Merck, 115930, Germany) in 1% sodium borate (Panreac, 141644, Spain).

The last semithin section (which corresponds to the section immediately adjacent to the block surface) was examined under light microscope and photographed to accurately locate the neuropil regions to be examined (Fig. 2). The blocks containing the embedded tissue were then glued onto a sample stub using conductive adhesive tabs (EMS 77825-09, Hatfield, PA, USA). All the surfaces of the block —except for the one to be studied (the top surface) — were covered with silver paint (EMS 12630, Hatfield, PA, USA) to prevent charging artifacts. The stubs with the mounted blocks were then placed into a sputter coater (Emitech K575X, Quorum Emitech, Ashford, Kent, UK) and the top surface was coated with a 10–20 nm thick layer of gold/palladium to facilitate charge dissipation.

**Figure 2.**
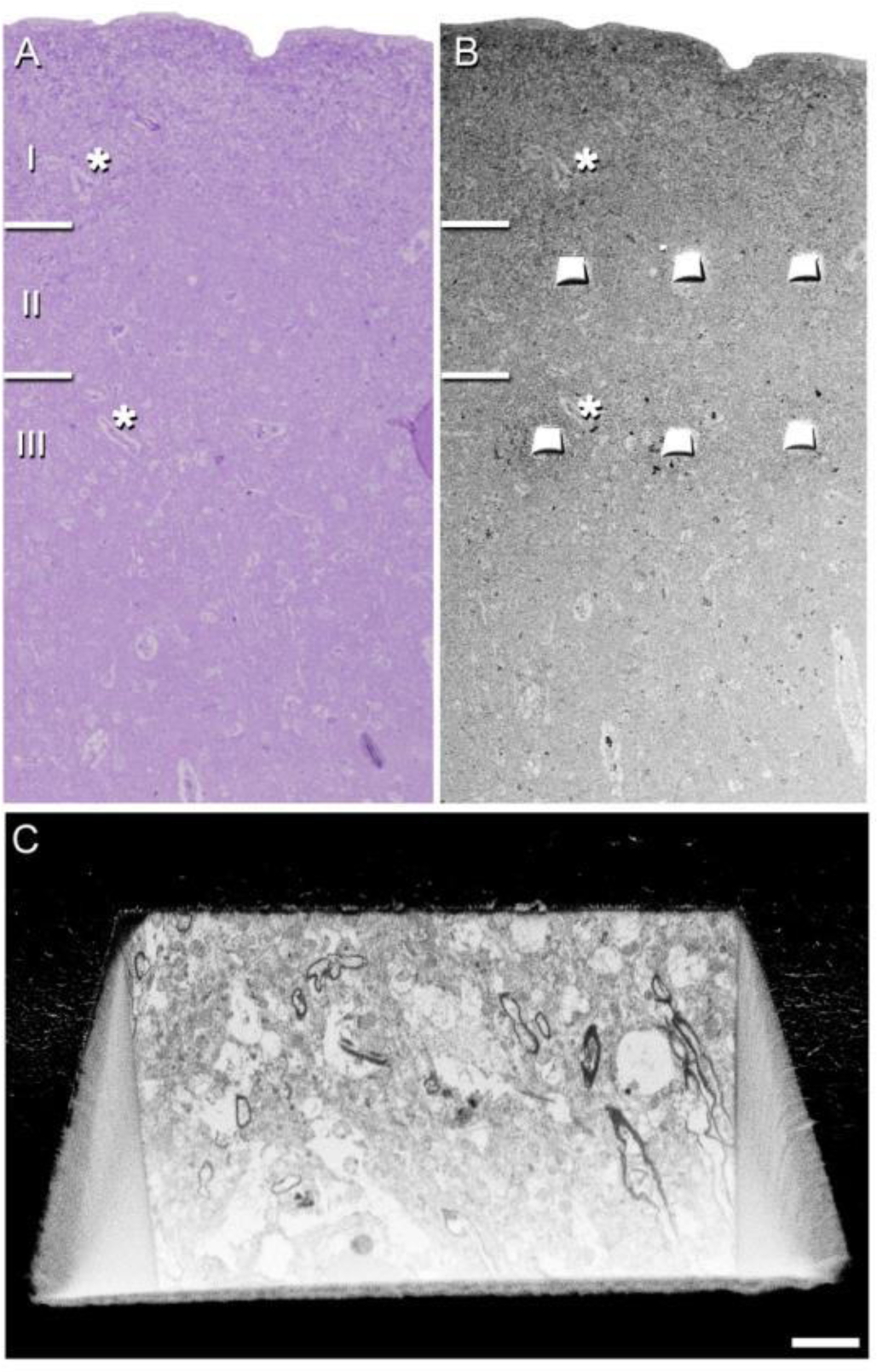
Correlative light/electron microscopy of layer II and III of the EC. Delimitation of layers is based on the staining pattern of 1 µm-thick semithin section, stained with toluidine blue (A), which is adjacent to the block for FIB/SEM imaging (B). (B) SEM image illustrating the block surface with trenches made in the neuropil (three per layer). Asterisks in A and B point to the same blood vessels, showing that the exact location of the region of interest was accurately determined. (C) SEM image showing the front of a trench made to expose the tissue and to acquire the FIB/SEM stack of images at the final magnification. Scale bar (in C): 70µm in A; 85µm in B; 3µm in C.

### Cortical thickness estimation

In order to estimate the atrophy of the EC, we measured the cortical thickness in three to five toluidine blue-stained semithin sections from each of the cases, obtained in the coronal plane of the cortex and containing the entire cortex, tracing a perpendicular line from the pial surface to the white matter. Measurements of the distance between the pial surface and the boundary with the white matter were performed using Fiji program (ImageJ 1.51; NIH, USA; http://imagej.nih.gov/ij/). To average data, three measurements were made per section.

### Volume fraction estimation of cortical elements

Three to five semithin sections (1–2μm thick) stained with 1% toluidine blue were used to estimate the respective volume fractions (Vv) occupied by blood vessels, cell bodies (glia and neurons) and neuropil. This estimation was performed applying the Cavalieri principle (Gundersen *et al*., 1988) by point counting using the integrated Stereo Investigator stereological package (Version 8.0, MicroBrightField Inc., VT, USA) attached to an Olympus light microscope (Olympus, Bellerup, Denmark) at 40x magnification. A grid, whose points covered an area of 400µm^2^, was overlaid over each semithin section to determine the V_v_ occupied by the different elements: neurons, glia, blood vessels and neuropil.

### Three-dimensional electron microscopy

The araldite block containing the tissue was used to obtain images stacks from the EC (Fig. 2) using a dual beam microscope (FIB/SEM; Crossbeam® 540 electron microscope, Carl Zeiss NTS GmbH, Oberkochen, Germany).

The dual beam microscope combines a high-resolution field-emission SEM column with a focused gallium ion beam (FIB), which permits removal of thin layers of material from the sample surface on a nanometer scale. As soon as one layer of material (20 nm thick) is removed by the FIB, the exposed surface of the sample is imaged by the SEM using a backscattered electron detector. The sequential automated use of FIB milling and SEM imaging allowed us to obtain long series of photographs of a 3D sample of selected regions (Merchán-Pérez et al., 2009). FIB/SEM images from the neuropil were obtained avoiding the neuronal and glial somata, blood vessels and also avoiding Aβ plaques to eliminate the effect of alterations of synapses in the vicinity of Aβ-plaques, which has been described previously (e.g., see Blazquez-Llorca et al., 2013).

In total, 24 stacks of images of the neuropil from layer II and III of the EC from AD cases were obtained (three stacks for each of the 4 cases, with a total volume studied of 9,006 μm^3^, Table 2). The data obtained from these stacks of images were compared to the data obtained in the same regions of four control cases (Domínguez-Álvaro *et al*., 2021).

**Table 2.**
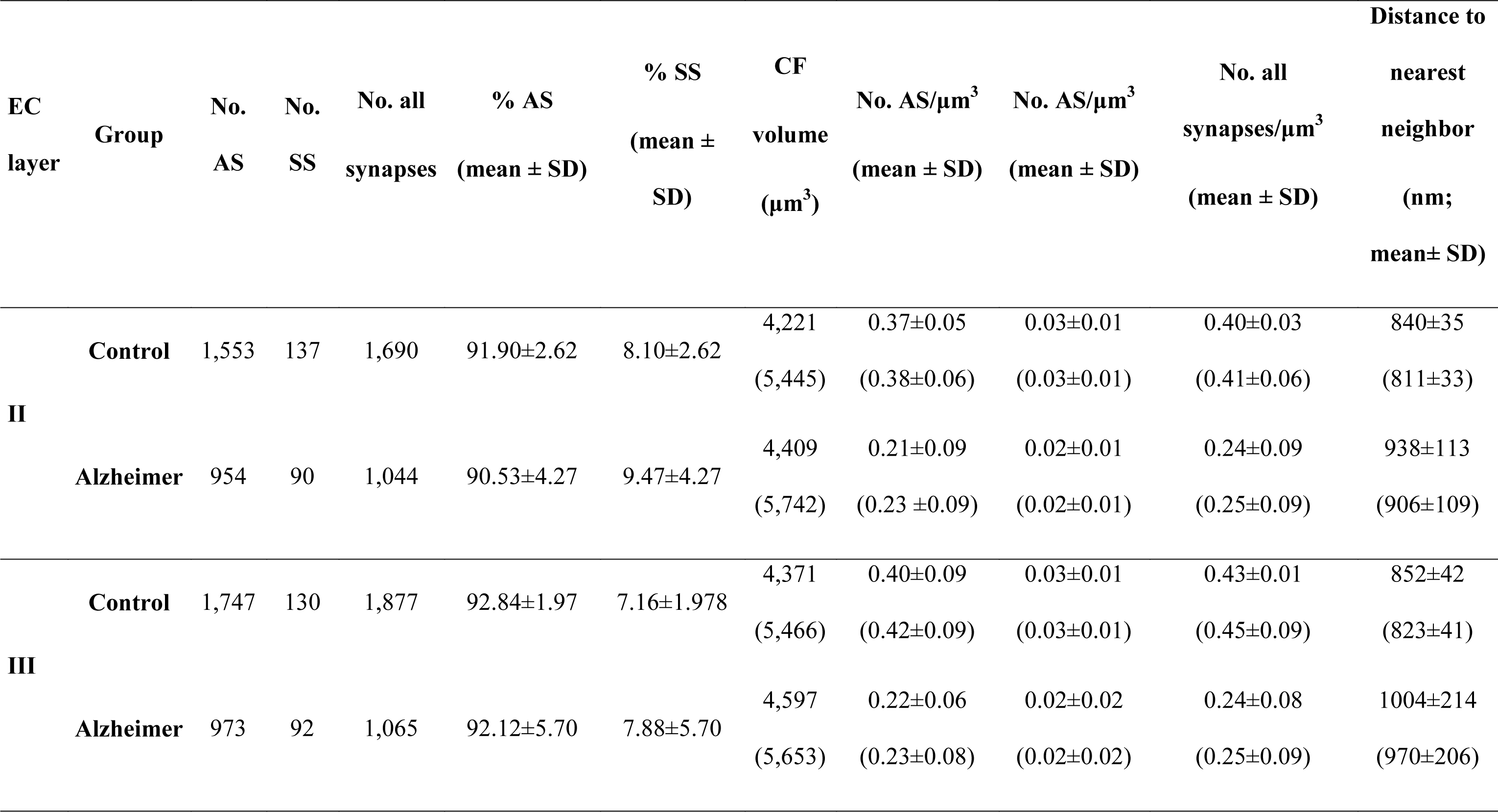
Accumulated data obtained from the ultrastructural analysis of the neuropil from layers II and III of the EC. All volume data are corrected for shrinkage factor. Data in parentheses are not corrected with the shrinkage and the fixation artifact factors. AS: asymmetric synapses; CF: counting frame; EC: entorhinal cortex; SD: standard deviation; SS: symmetric synapses. The data for individual cases are shown in Extended Table 2-1.

### Synaptic three-dimensional analysis

FIB/SEM stacks of images were analyzed using EspINA software (*EspINA Interactive Neuron Analyzer*, 2.1.9; https://cajalbbp.es/espina/), which allows the segmentation of synapses in the reconstructed 3D volume (Morales *et al*., 2011; Fig. 3). Since the EspINA software allows navigation through the stack of images (Figs. 3, 4; Movies 1, 2), it was possible to unambiguously identify every synapse as asymmetric synapses (AS) or symmetric synapses (SS), based on the thickness of the PSD (Fig. 4). There is a consensus for classifying cortical synapses into asymmetric synapses (AS; or type I) and symmetric synapses (SS; or type II). The main characteristic distinguishing these synapses is the prominent or thin post-synaptic density, respectively: synapses with prominent PSDs are classified as AS, while thin PSDs are classified as SS (Gray, 1959; Peters and Palay, 1991).

**Figure 3.**
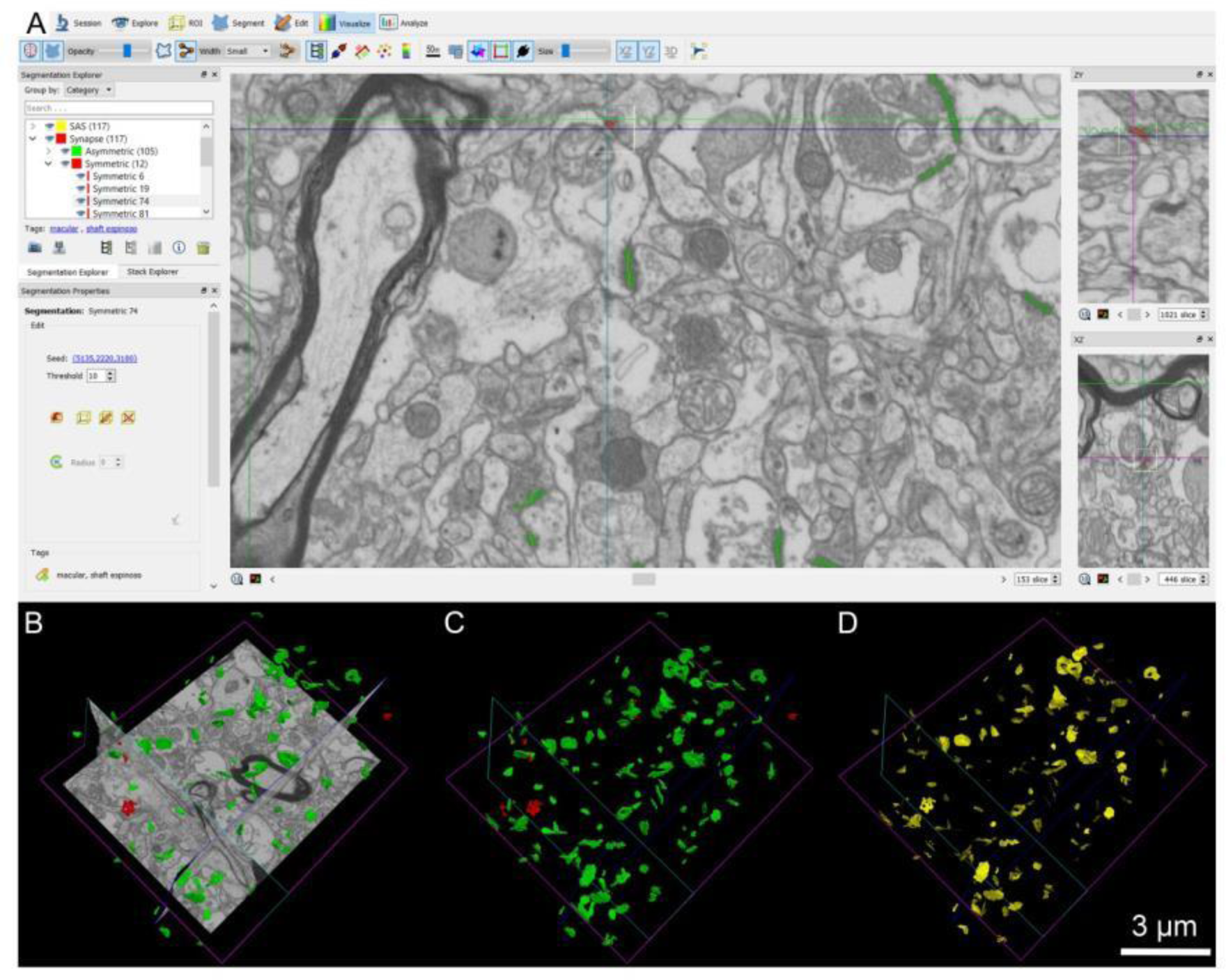
Screenshot of the EspINA software user interface. **(A)** In the main window, the sections are viewed through the xy plane (as obtained by FIB/SEM microscopy). The other two orthogonal planes, yz and xz, are also shown in adjacent windows on the right. **(B)** The 3D windows show the three orthogonal planes and the 3D reconstruction of AS (green) and SS (red) segmented synapses, the reconstructed synapses **(C)**, and the computed SAS for each reconstructed synapse (in yellow; **D**).

**Figure 4.**
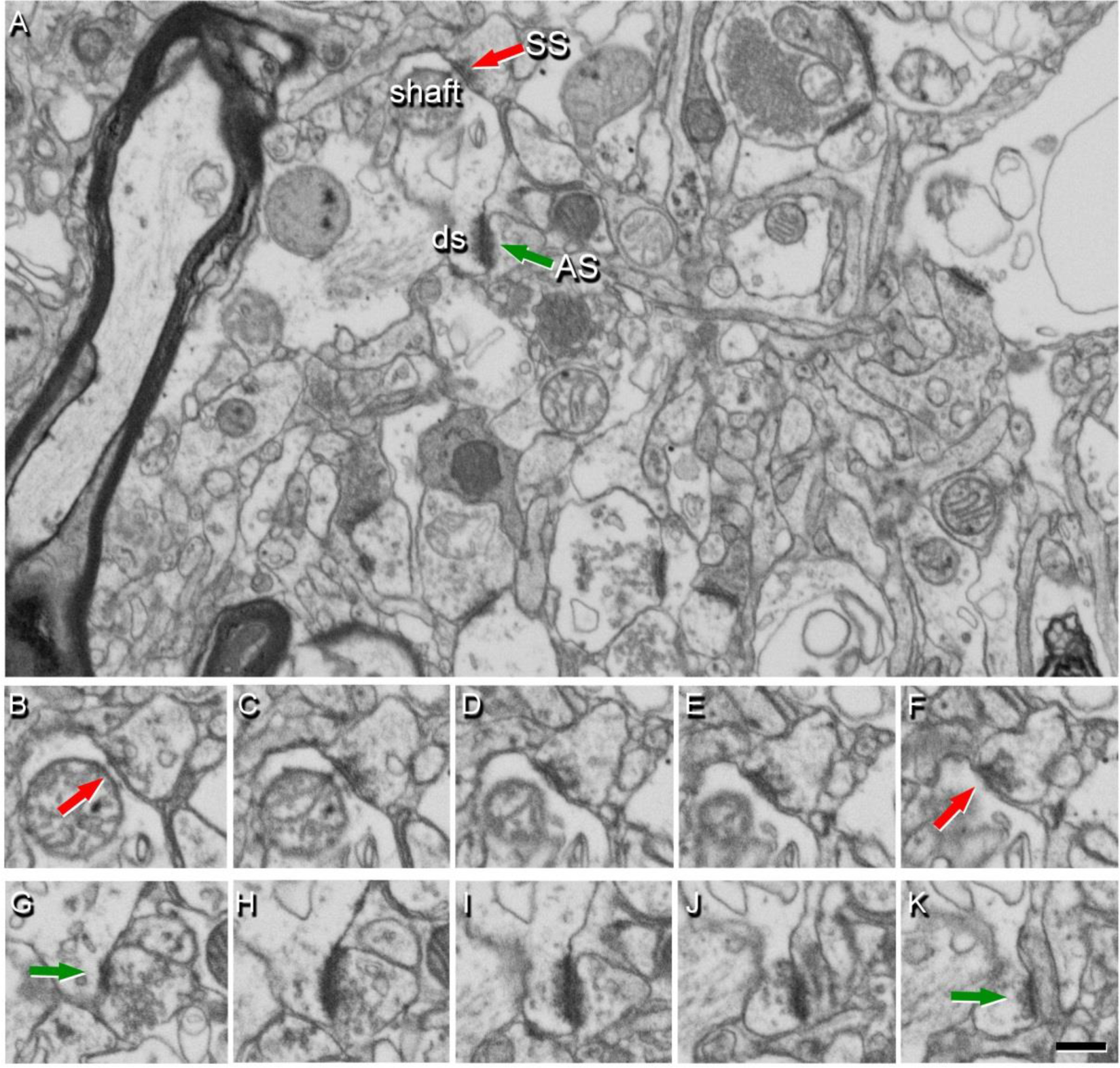
Identification of synapses in serial electron microscopy images obtained by FIB/SEM. **(A)** Image from layer II showing the neuropil from an AD case, with two synapses indicated (arrows) as examples of symmetric synapse (SS, red) on a dendritic shaft, and asymmetric synapse (AS, green) on a dendritic spine head (ds). Synapse classification was based on examination of the full sequence of serial images; the SS can be visualized in **B−F**, and the AS in **G−K**. Scale bar (in K): 1,000 nm in A; 1,300 nm in B−K.

EspINA also allowed the application of an unbiased 3D counting frame (CF) to obtain the synaptic density per volume (for details, see Merchán-Pérez *et al*., 2009). The synaptic density values were obtained by dividing the total number of synapses by the total volume of the CF. Geometrical characteristics —such as size— and spatial distribution features (centroids) of each reconstructed synapse were also calculated by EspINA.

EspINA software extracts the Synaptic Apposition Area (SAS) and provides its morphological measurements (Fig. 3). Since the pre- and post-synaptic densities are located face to face, their surface areas are comparable (for details, see Morales et al., 2013). Since the SAS comprises both the active zone and the PSD, it is a functionally relevant measure of the size of a synapse (Morales et al., 2013). In addition, the visualization of each 3D reconstructed synapse allowed us to determine the synaptic morphology. Based on the presence of perforations or indentations in their perimeters, the synapses could be classified into four types: macular (with a flat, disk-shaped PSD), perforated (with one or more holes in the PSD), horseshoe (with an indentation in the perimeter of the PSD) or fragmented (with two or more physically discontinuous PSDs) (for a detailed description, see Domínguez-Álvaro *et al*., 2019).

To identify the postsynaptic targets of the synapses, we navigated the image stack using EspINA to determine whether the postsynaptic element was a dendritic spine (‘spine’ or ‘spines’, for simplicity) or a dendritic shaft (Fig. 5; Movies 1, 2). Unambiguous identification of spines as postsynaptic targets requires the spine to be visually traced to the parent dendrite, yielding what we refer to as fully reconstructed spines. Additionally, when synapses were established on a spine head-shaped postsynaptic element whose neck could not be followed to the parent dendrite, or whose neck was truncated, we identified these elements as non-fully reconstructed spines. These non-fully reconstructed spines were identified on the basis of their size and shape, the lack of mitochondria and the presence of a spine apparatus — or because they were filled with a characteristic fluffy material (used to describe the fine and indistinct filaments present in the spines) — a term coined by Peters et al. (1991) (see also del Río and DeFelipe, 1995). For simplicity, we will refer to the fully reconstructed and non-fully reconstructed spines as spines, unless otherwise specified. Similarly, for the unambiguous identification of dendritic shafts, it is necessary to be able to visually trace them inside the stack. Accordingly, when the postsynaptic element of a synapse was close to the margins and it was truncated by the borders of the stack, the identity of the postsynaptic target could not be determined. Therefore, the targets of synapses in each of the stacks were classified into two main categories: spines and dendritic shafts, while truncated elements that could not be safely identified were discarded. When the postsynaptic target was a spine, we further recorded the position of the synapse on the head or neck. Additionally, when the postsynaptic element was identified as a dendritic shaft, it was classified as “with spines” or “without spines”.

**Figure 5.**
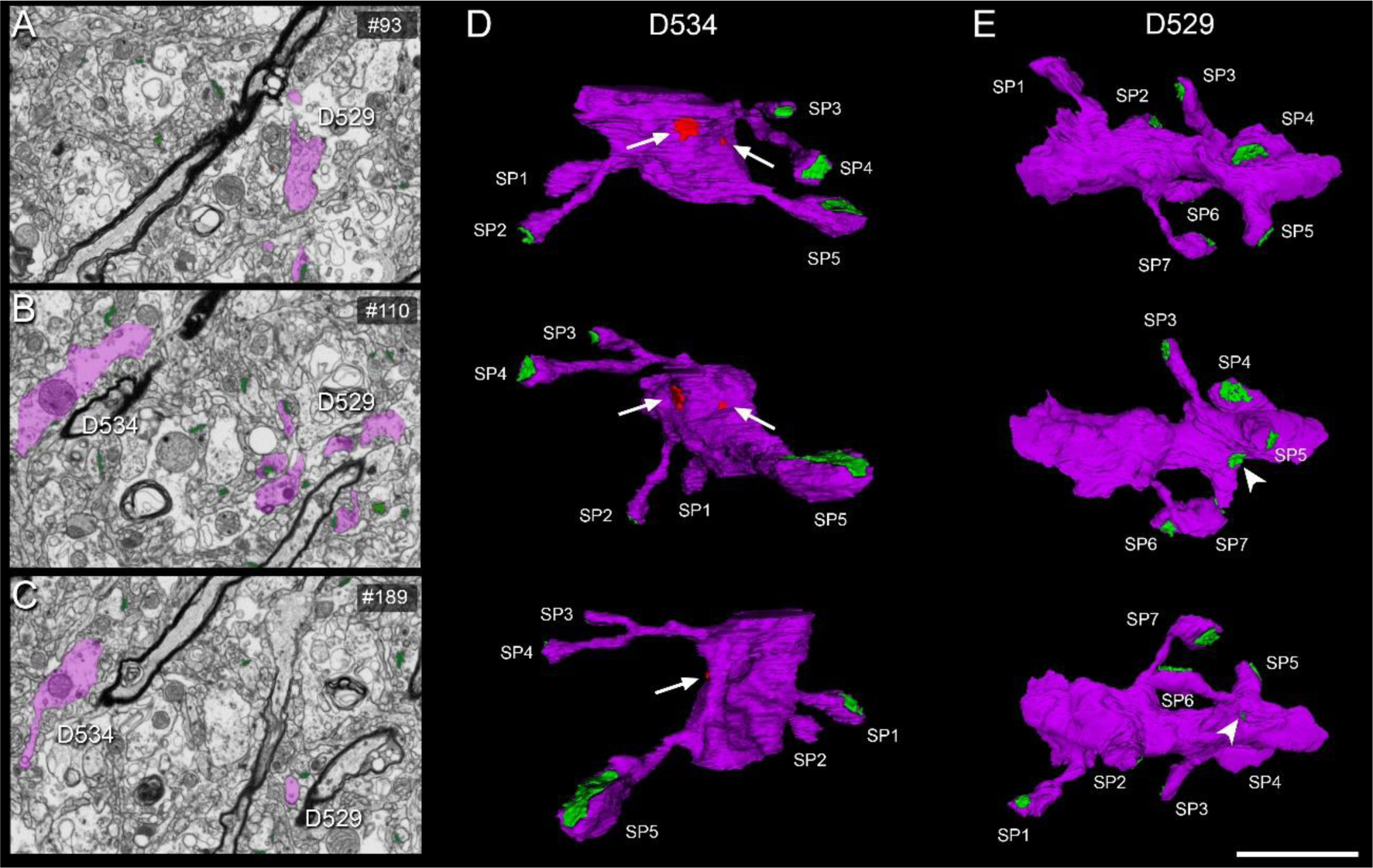
3D reconstruction of dendritic segments from FIB/SEM serial images. (A–C) Serial images showing two dendritic segments (D529, D534) partially reconstructed (in purple). 3D Reconstructions of these dendritic segments are displayed in panels D and E. Panel D shows the 3D reconstructed dendritic segment D534 in three views after rotation about the major dendritic axis. Five dendritic spines (SP 1–5) are shown, establishing asymmetric synapses (green) — and two symmetric synapses (arrows) on the shaft (red) are also visible. Panel E shows the 3D reconstructed dendritic segment D534 in three views after rotation about the major dendritic axis. Seven dendritic spines (SP 1–7) establishing asymmetric synapses (green) are illustrated. Note that the synapse on SP4 can be identified as perforated in the middle panel (E). The arrowhead indicates an asymmetric synapse on a spine neck. Scale bar (in E) indicates 3 µm in A–C and 2 µm in D, E.

### Spatial Distribution Analysis of Synapses

To analyze the spatial distribution of synapses, spatial point-pattern analysis was performed as described elsewhere (Merchán-Pérez *et al*., 2014). For each of the 24 different samples, we calculated three functions commonly used for spatial point pattern analysis: F, G and K functions (for a detailed description, see Blazquez-Llorca *et al*., 2015). Additionally, we measured the distance of each synapse to its nearest synapse, to compare control and AD samples. This study was carried out using the Spatstat package and R Project program (Baddeley *et al*., 2015).

### Tissue shrinkage

All measurements were corrected for the tissue shrinkage that occurs during osmication and plastic-embedding of the vibratome sections containing the area of interest, as described by (Merchán-Pérez et al., 2009). We measured the surface area and thickness of the vibratome sections with Stereo Investigator (MBF Bioscience, Williston, VT, USA), both before and after they were processed for EM (Oorschot et al., 1991). The surface area after processing was divided by the value before processing to obtain an area shrinkage factor (p2) of 0.933. The linear shrinkage factor for measurements in the plane of the section (p) was therefore 0.966. The shrinkage factor in the z-axis was 0.901. In addition, the total volume was corrected for the presence of fixation artifacts, which did not affect the accurate identification and quantitation of synapses (i.e., swollen neuronal or glial processes). The volume occupied by these artifacts was calculated applying the Cavalieri principle (Gundersen et al., 1988) and was discounted from the volume of the stacks of images to avoid underestimation of the number of synapses per volume. Specifically, a stereological grid with an associated area per point of 400,000 nm2 was superimposed onto each FIB/SEM stack using Image J Stereology Toolset (Mironov, 2017). Estimations were made every 20th section in each of the stacks. Every FIB/SEM stack was examined and the volume artifact ranged from 1 to 16% of the volume stacks. Volume fraction estimation was performed by point counting using the Cavalieri principle (Gundersen et al., 1988), in a similar fashion to the volume fraction estimation of cortical elements in semithin sections (see “Volume fraction estimation of cortical elements”).

All parameters measured were corrected to obtain an estimate of the pre-processing values. The shrinkage factor was used to correct the synaptic apposition surface (SAS) area and perimeter data, while both the shrinkage and the fixation artifact factors were used to correct synaptic density values.

### Statistical analysis

To determine possible differences between the control and AD samples, statistical comparisons of the following parameters were carried out using the unpaired Mann-Whitney (MW) nonparametric U-test: cortical thickness; synaptic density; type of synapse (AS or SS); SAS size; synaptic shape; postsynaptic targets; and distance to the nearest neighboring synapse. To determine possible differences between control and AD samples regarding volume fractions (neuronal and glial cell bodies, blood vessels and neuropil), parametric t-test were carried out. Frequency distribution analyses were performed using Kolmogorov-Smirnov (KS) nonparametric test.

To perform statistical comparisons of AS and SS proportions, chi-square (χ²) test was used for contingency tables. The same method was used to study whether there were significant differences between groups in relation to the shape of the synaptic junctions and their postsynaptic target. In all the χ² statistical analyses, we firstly performed an “Omnibus test” based on 2x4 contingency tables. To further investigate the specific cells driving the significance of the χ² test, a partitioning procedure was applied to create 2x2 contingency tables (Sharpe, 2015).

Statistical analyses were performed using the GraphPad Prism statistical package (Prism 8.4.2 for Windows, GraphPad Software Inc., USA) and SPSS (IBM SPSS Statistics v24, IBM Corp., Armonk, NY, USA).

## Results

Data regarding the synaptic organization of the human layers II and III from the EC in the four control subjects has been previously published and detailed information can be found therein (Domínguez-Álvaro *et al*., 2021). What follows are the alterations of synapses in AD cases in comparison with control cases.

### Cortical thickness

Measurements of the cortical thickness were derived from semi-thin toludine blue (serial sections) using light microscopy. The mean cortical thickness of the EC was 2.0±0.17 mm (mean±SD) in the control group and 1.6±0.49 mm in the cases with AD. Although approximately 20% lower EC cortical thickness was observed in AD cases was observed, we did not find a statistically significant difference between the two groups (MW, p = 0.34; Extended Table 1-1).

### Volume fraction of cortical elements

In the control group layer II, the values of the estimated volume fraction (Vv) occupied by neuronal somata, glial somata, blood vessels and neuropil were 4.5%, 2.6%, 3.8% and 89.1%, respectively. In the group with AD, these values were 2.6%, 4.6%, 2.3% and 90.5%, respectively. The only statistically significant difference was that of the Vv of glial somata, which was found to be significantly higher in cases with AD (t-test, p=0.03) (Fig. 6; Extended Tables 1-2, 1-3).

**Figure 6.**
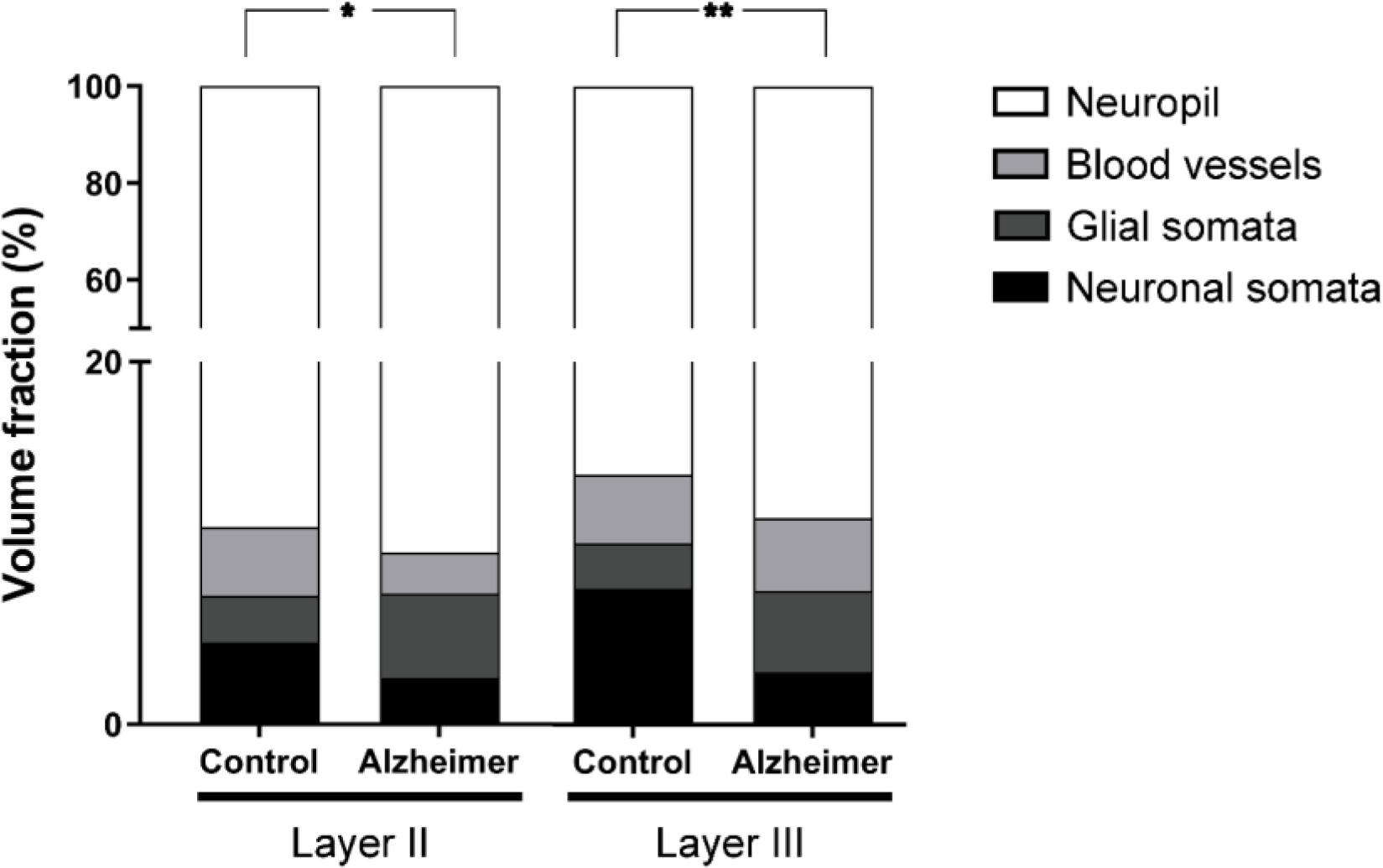
Graph showing the volume fraction (Vv) occupied by neuronal and glial somata, blood vessels and neuropil in both layers II and III of the EC from control subjects and AD cases. Asterisks show the differences between groups. In layer II, a significantly higher Vv of glial somata was found in AD cases (t-test, p = 0.03). In layer III, a significantly lower Vv of neuronal somata was found in AD patients (t-test, p = 0.007). Mean values and case values are detailed in Extended Tables 1-2 and 1-3.

**Table 3.** Area (mean ± sem, in nm^2^) and perimeter (mean ± sem, in nm) of the SAS in layers II and III of the EC. Data on area and perimeter are corrected for shrinkage factor (Data in parentheses are not corrected with the shrinkage and the fixation artifact factors). AS: asymmetric synapses; EC: entorhinal cortex; SAS: synaptic apposition surface; sem: standard error of the mean; SS: symmetric synapses. The data for individual cases are shown in Extended Table 3-1.

| EC layer | Group | Type of synapse<br>(no. of synapses) | Area of SAS | Perimeter of SAS |
| --- | --- | --- | --- | --- |
| II | Control | AS | 110,311 ±3,229 | 1,631 ±47 |
|  |  | (1552) | (102,865) | (1,575) |
| | | SS | 65,997 ±6,041 | 1,404 ±73 |
|  |  | (139) | (61,543) | (1,356) |
| | Alzheimer | AS | 136,166 ±10,773 | 1,819 ±52 |
|  |  | (952) | (126,976) | (1,757) |
| | | SS | 59,232 ±6,589 | 1,239 ±72 |
|  |  | (90) | (55,234) | (1,196) |
| III | Control | AS | 124,183 ±3,265 | 1,826 ±67 |
|  |  | (1746) | (115,801) | (1,763) |
| | | SS | 67,445 ±3,243 | 1,399 ±75 |
|  |  | (129) | (62,893) | (1,351) |
| | Alzheimer | AS | 122,727 ±10,453 | 1,720 ±116 |
|  |  | (973) | (114,444) | (1,661) |
| | | SS | 57,271 ±14,132 | 1,343 ±272 |
|  |  | (94) | (56,082) | (1,348) |

In layer III of the control group, the Vv of neuronal somata, glial somata, blood vessels and neuropil were 7.5%, 2.5%, 3.8% and 86.2%. In cases with AD, these values were 2.9%, 4.5%, 4.0% and 88.6%. Statistical comparisons showed a significantly lower Vv of neuronal somata (t-test, p=0.007) in AD (Fig. 6; Extended Tables 1-2, 1-3).

### Synaptic Density and AS:SS ratio

A total of 2,019 synapses from AD cases (1,044 synapses in layer II and 1,065 in layer III) were identified and reconstructed in 3D, after discarding incomplete synapses or those touching the exclusion edges of the CF, and the total volume analyzed was 9,006 μm^3^ in AD samples (Table 2 and Extended Table 2-1).

A significantly lower synaptic density (taking both AS and SS together) was observed in AD cases — this value was 40% lower than in control cases in both EC layers. In layer II, we found 0.24 synapses/μm^3^ in AD cases and 0.40 synapses/μm^3^ in control cases (MW, p=0.03; Fig. 7A). In layer III, the mean synaptic density in AD was 0.24 synapses/μm^3^ compared to 0.43 synapses/μm^3^ found in control cases (MW, p=0.03; Fig. 7B; Table 2 and Extended Table 2-1).

**Figure 7.**
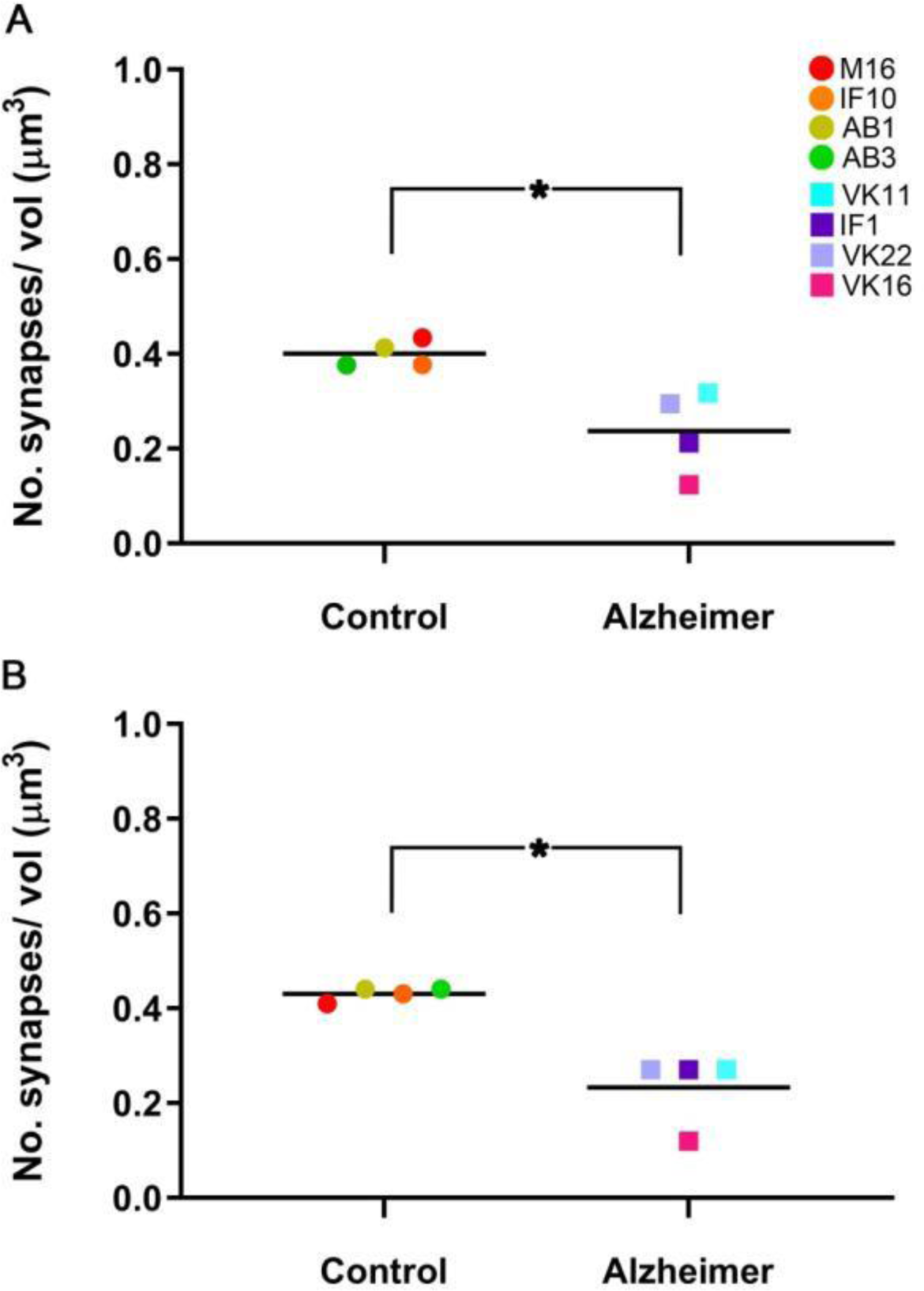
Graphs showing the overall mean synaptic density in layer II (A) and layer III (B) of the EC in control and AD cases. Control cases are represented by circles and AD cases are represented by squares. Each color corresponds to each case analyzed, as denoted in the upper right-hand corner. Asterisks show significant differences between groups: in both layer II and layer III of the EC, the mean synaptic density was significantly lower in AD patients (MW, p = 0.03).

In layer II and III, the AS:SS ratio for the AD cases was very similar to that of the control cases (close to 92:8 in both layers and cases; Table 2 and Extended Table 2-1), and no statistically significant differences were found between the control group and the AD group in any layer (χ², p>0.05).

### Three-dimensional spatial synaptic distribution

Analysis indicates a clear fit to a complete spatial randomness (CSR) model, since F, G and K functions closely resemble the theoretical curve that these functions represent, both in the control and the AD cases. The CSR model defines a situation where a point is equally likely to occur at any location within the study volume, regardless of the locations of other points. Therefore, a CSR model is considered as a reference for a random pattern in spatial point process statistics (reviewed in Merchán-Pérez et al., 2014). That is, the spatial distribution of the synapses fitted a random distribution in all subjects and layers. Furthermore, the estimation of the distance from each synapse to its closest synapse showed that although the mean distance was greater in AD cases in both layer II (811 nm in control; 906 nm in AD; Table 2 and Extended Table 2-1) and layer III (823 nm in control, 970 nm in AD; Table 2 and Extended Table 2-1), no statistically significant differences were found (MW, p>0.05).

### Study of the synapse characteristics

#### Synaptic size

The AD samples did not show differences between layers in the SAS area of AS (MW, p=0.41; Table 3 and Extended Table 3-1). However, we found significantly larger mean values of SAS area for AS (MW, p=0.03) from layer II in AD cases compared to the controls. These differences were not found in SS or in layer III (MW, p>0.05; Table 3 and Extended Table 3-1).

Analysis of the frequency distributions of the area and perimeter of the AS showed significant differences in the frequency distributions of area in layer II and in the perimeter in layer III (KS, p<0.0001), indicating a lower proportion of small SAS area in layer II in AD cases and higher proportions of small SAS perimeter in layer III in AD cases (Fig. 8).

**Figure 8.**
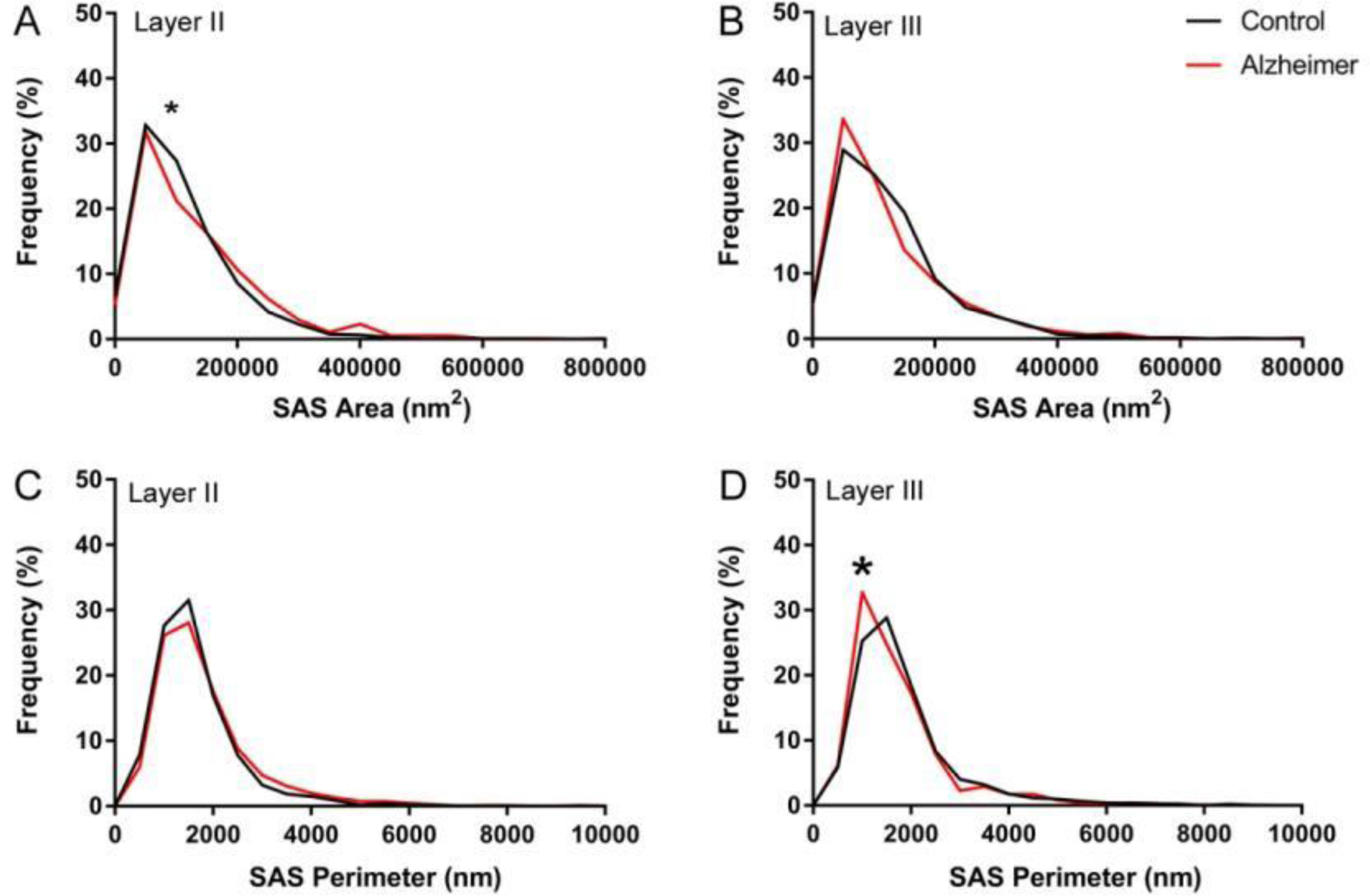
Graph showing the frequency distribution plots of AS SAS area (A, B) and perimeter (C, D), in layers II and III of the EC in control subjects (black) and AD patients (red). Statistical comparisons between groups showed significant differences (KS, p<0.0001; indicated with an asterisk) in the frequency distributions of SAS area in layer II **(A)**, and in the frequency distribution of SAS perimeter in layer III **(D)**.

#### Synaptic shape

Most synapses, AS and SS, presented a macular shape in both control and AD cases in layers II and III (>80%), while synapses with more complex shapes (i.e., horseshoe-shaped, perforated or fragmented) were less frequent (Fig. 9; Table 4, Extended Tables 4-1 and 4-2). Evaluation of the possible differences between the control and the AD group in layer II showed a slightly higher proportion of horseshoe-shaped AS in AD samples (χ², p = 0.04; Table 4 and Extended Table 4-1; Fig. 9B). Analysis of layer III revealed that fragmented (χ^2^, p=0.0004) and macular (χ^2^, p<0.0001) AS were more frequent in the AD group, whereas perforated AS were less frequent in AD cases than in the control individuals (χ², p <0.0001; Table 4 and Extended Table 4-2; Fig. 9C). Analysis of the SS in layers II and III did not reveal differences between groups (χ², p>0.001).

**Figure 9.**
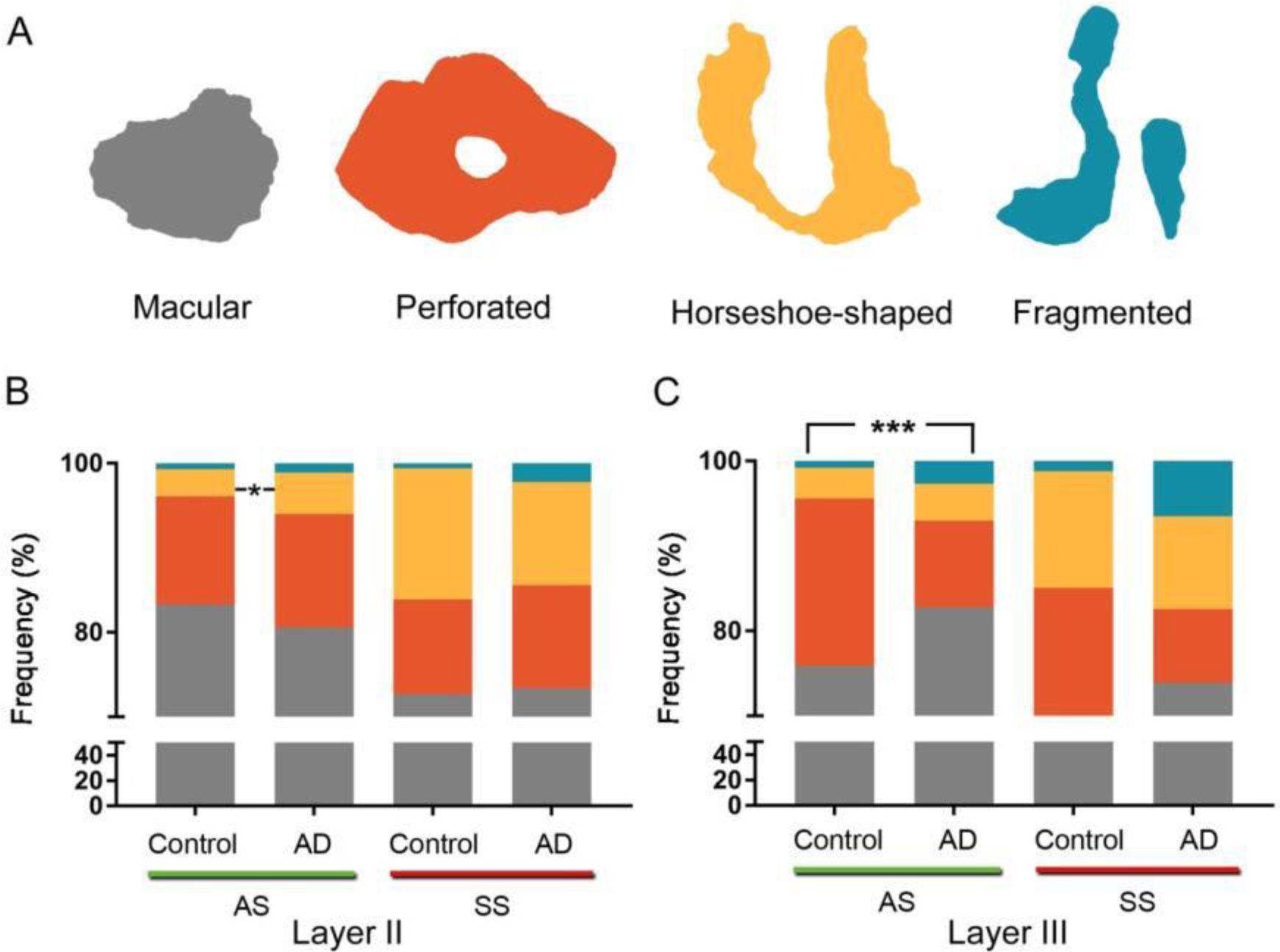
Proportions of the different synaptic shapes (A) in layers II (B) and III (C) of the EC in control subjects and AD patients. **(A)** Schematic representation of the synaptic shapes: macular synapses, with a continuous disk-shaped PSD; perforated synapses, with holes in the PSD; horseshoe-shaped, with a tortuous horseshoe-shaped perimeter with an indentation; and fragmented synapses, with two PSDs with no connections between them. Proportions of macular, perforated, horseshoe-shaped and fragmented AS and SS are displayed for layer II **(B)** and III **(C)** of the EC in control subjects and AD cases. Statistical differences (indicated with asterisks) showed that in layer II, horseshoe-shaped AS were more frequent (χ^2^, p=0.04) in AD cases; and in layer III of the AD group, fragmented (χ^2^, p=0.0004) and macular (χ^2^, p<0.0001) AS were more frequent, whereas perforated AS were less frequent in AD cases than in control individuals (χ², p <0.0001).

**Table 4.** Proportion of the different synaptic shapes in layers II and III of the EC. Data are given as percentages with the absolute number of synapses studied in parentheses. AS: asymmetric synapses; EC: entorhinal cortex; SS: symmetric synapses. Data for individual cases are shown in Extended Tables 4-1 and 4-2.

| **EC layer** | **Group** | **Type of synapse** | **Macular synapses** | **Perforated synapses** | **Horseshoe synapses** | **Fragmented synapses** | **Total synapses** |
| --- | --- | --- | --- | --- | --- | --- | --- |
| **II** | **Control** | AS | 83.0% | 12.9% | 3.3% | 0.8% | 100% |
|  |  |  | (1,285) | (200) | (51) | (13) | (1,549) |
|  |  | SS | 73.0% | 11.0% | 15.3% | 0.7% | 100% |
|  |  |  | (100) | (15) | (21) | (1) | (137) |
|  | **Alzheimer** | AS | 80.6% | 13.4% | 4.9% | 1.1% | 100% |
|  |  |  | (769) | (128) | (47) | (10) | (954) |
|  |  | SS | 73.4% | 12.2% | 12.2% | 2.2% | 100% |
|  |  |  | (66) | (11) | (11) | (2) | (90) |
| **III** | **Control** | AS | 75.9% | 19.9% | 3.3% | 0.9% | 100% |
|  |  |  | (1,326) | (347) | (58) | (15) | (1,746) |
|  |  | SS | 65.9% | 15.5% | 17.0% | 1.6% | 100% |
|  |  |  | (85) | (20) | (22) | (2) | (129) |
|  | **Alzheimer** | AS | 82.8% | 10.2% | 4.3% | 2.7% | 100% |
|  |  |  | (806) | (99) | (42) | (26) | (973) |
|  |  | SS | 73.9% | 8.7% | 10.9% | 6.5% | 100% |
|  |  |  | (68) | (8) | (10) | (6) | (92) |

#### Analysis of Synaptic Size and Shape

Additionally, we determined whether the shape of the synapses was related to their size. For this purpose, the area and perimeter of the SAS of AS were analyzed according to their synaptic shape (SS were not analyzed since the sample size was too small).

When we compared the synaptic size considering the synaptic shape, we found that the mean SAS area from perforated AS in layer III was larger in AD samples (296,156±24,116 nm^2^, mean±sem) than in the controls (228,057±6,993 nm^2^, mean±sem; MW, p=0.03; Extended Tables 4-3, 4-4). No differences were found in the mean area and perimeter of the SAS from macular, horseshoe-shaped and fragmented synapses (MW, p>0.05). Analysis of the frequency distribution of the area and perimeter showed no differences between groups (Extended Fig. 9-1; KS; p>0.0001).

#### Study of the postsynaptic targets

The postsynaptic targets of 1,706 synapses were determined from the AD samples. In layer II, we identified the postsynaptic targets of a total of 807 AS and 82 SS (Table 5 and Extended Table 5-1), and in layer III, the postsynaptic targets for 732 AS and 85 SS were determined (Table 5 and Extended Table 5-2).

**Table 5.** Distribution of asymmetric (AS) and symmetric (SS) synapses on spines and dendritic shafts in layers II and III of the EC. Synapses on spines have been sub-divided into those that are established on spine heads and those that are established on spine necks. Moreover, we differentiated between aspiny and spiny dendritic shafts. Data are expressed as percentages with the absolute number of synapses studied given in parentheses. Data for each individual case are shown in Extended Tables 5-1 and 5-2. AD: Alzheimer’s disease; AS: asymmetric synapses; EC: entorhinal cortex; SS: symmetric synapses.

| Layer | Group | Type of syn. | On spine heads | On nf. spines | On spine necks | On aspiny dendritic shaft | On spiny dendritic shaft | Total |
| --- | --- | --- | --- | --- | --- | --- | --- | --- |
| II | C | AS | 39.68%<br>(567) | 18.89%<br>(270) | 0.56%<br>(8) | 19.87%<br>(284) | 20.99%<br>(300) | 100%<br>(1,429) |
|  |  | SS | 9.38%<br>(12) | 4.69%<br>(6) | 0.78 %<br>(1) | 22.66%<br>(29) | 62.50%<br>(80) | 100%<br>(128) |
|  | AD | AS | 35.19%<br>(284) | 17.84%<br>(144) | 0.87%<br>(7) | 25.15%<br>(203) | 20.94%<br>(169) | 100%<br>(807) |
|  |  | SS | 10.98%<br>(9) | 1.22%<br>(1) | 0% (0) | 48.78%<br>(40) | 39.02%<br>(32) | 100%<br>(82) |
| III | C | AS | 39.68%<br>(642) | 15.33%<br>(248) | 0.74%<br>(12) | 17.49%<br>(283) | 26.79%<br>(433) | 100%<br>(1,618) |
|  |  | SS | 11.90%<br>(15) | 4.76%<br>(6) | 2.38%<br>(3) | 22.22%<br>(28) | 58.73%<br>(74) | 100%<br>(126) |
|  | AD | AS | 41.67%<br>(305) | 11.75%<br>(86) | 0.82%<br>(6) | 28.83%<br>(211) | 16.94%<br>(124) | 100%<br>(732) |
|  |  | SS | 5.88%<br>(5) | 1.18%<br>(1) | 3.53%<br>(3) | 40% (34) | 49.41%<br>(42) | 100%<br>(85) |

The most abundant type of synapse in AD individuals was AS on spine heads (range 47.9-48.1%) closely followed by AS on dendritic shafts (41−41.8%). SS on dendritic shafts were less common (8.1−9.3%), SS on spine heads were very uncommon (0.7-1.1%), and AS or SS on spine necks were very infrequent indeed (<1%; Fig. 10).

**Figure 10.**
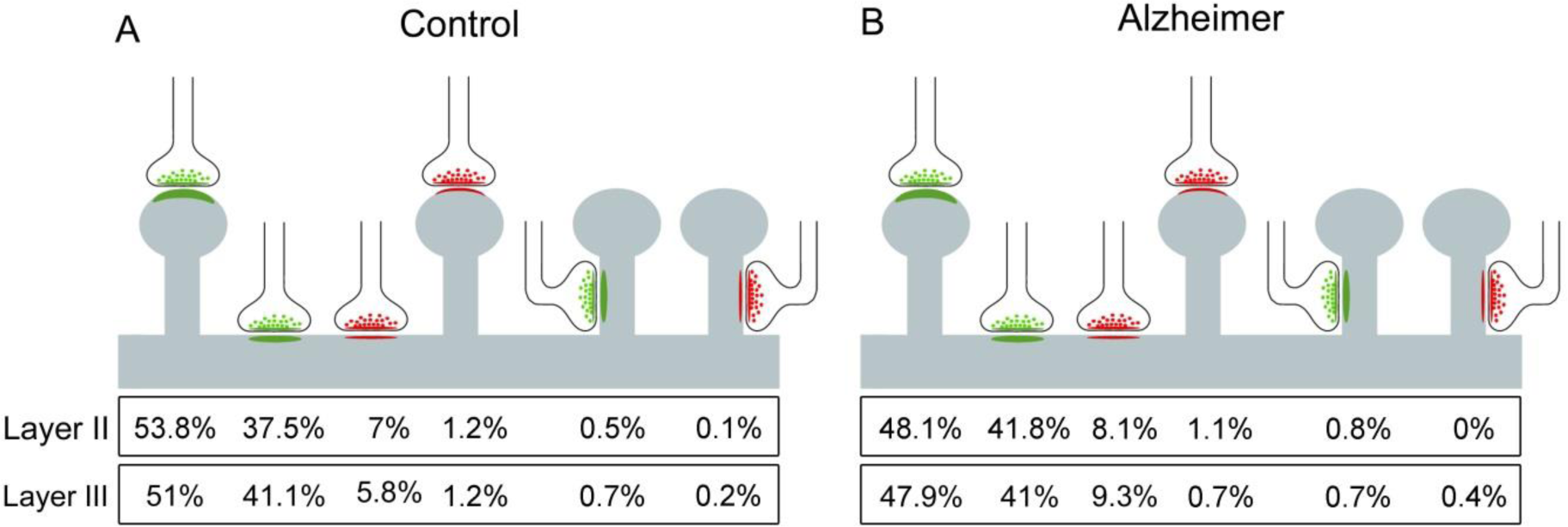
Postsynaptic target distribution in layers II and III of the EC. Schematic representation of the distribution of AS and SS on different postsynaptic targets from control cases **(A)** and AD patients **(B).** In both layer II and layer III, the AS preferentially targeted spine heads, while the SS preferentially targeted dendritic shafts. Percentages of postsynaptic targets are indicated, showing —from left to right— the most frequent type (AS on spine heads) to the least frequent type (SS on spine necks). Synapses on spines have been sub-classified into those that are established on the spine head and those established on the neck.

When the preference of the synaptic types (AS or SS) for a particular postsynaptic element was analyzed, we found that, in both controls and AD individuals, excitatory contacts (AS) presented a statistically significant preference for spines, whereas inhibitory contacts (SS) preferentially targeted dendritic shafts (χ^2^, p<0.001), in both layers, in line with the findings in control EC samples from the same layers (Domínguez-Álvaro *et al*., 2021). Contingency tables were used to assess possible differences in the distribution of postsynaptic elements of cases with AD. No statistically significant differences were found (χ², p>0.001), although a slightly lower proportion of AS established on spine heads (χ², p=0.02) and a higher proportion of AS on dendritic shafts were observed in AD samples from layer II (χ², p=0.03).

We also analyzed whether there was a relationship between the synapse size and the type of postsynaptic element. When we compared the synaptic size of AS associated with the postsynaptic targets between control cases and AD cases, no differences were found in any layer with regard to the area and perimeter of the SAS (MW, p>0.05) or their frequency distribution (KS, p>0.001; Extended Fig. 10-1).

## Discussion

In total, there are four main findings in the present study on the human EC: at the light microscope level, (i) a significantly lower volume fraction occupied by neuronal bodies in the layer III was found in AD cases; at the ultrastructural level, (ii) a significantly lower synaptic density was found in both layers in AD cases; (iii) synaptic morphology analysis revealed larger and more complex synapses in layer II in AD cases; and (iv) there was a greater proportion of small and simple synapses in layer III in AD cases than in control individuals.

### Cortical thickness and volume fraction analysis

At the light microscopic level, atrophy of the EC and adjacent cortical regions is clearly visible to the naked eye (Fig. 1). However, the difference in the mean cortical thickness was no evident (Extended Table 1-1). Macroscopic atrophy of the EC has been described in AD cases (Van Hoesen *et al*., 1991). This cortical atrophy and, more specifically, the neuronal loss (see below) observed in AD cases have been related to the presence of neurofibrillary tangles (NFTs) and Aβ plaques, with reports indicating an inverse relationship between the number of neurons and the degree of neuropathology (Gómez-Isla *et al*., 1996; Van Hoesen *et al*., 1991).

In AD brain samples, we found a significant lower neuronal volume fraction in the layer III. This is in line with the reported neuronal loss from AD cases in the EC (Gómez-Isla *et al*., 1996; Šimić *et al*., 1997; Van Hoesen and Hyman, 1990). Since the neurons of EC layer III constitute the projection elements of the perforant pathway to the hippocampal CA1 (together with neurons from layer II), this neuronal loss may contribute to the memory impairment described in AD cases (Andrade-Moraes et al., 2013, Gunten *et al*., 2006; Hyman *et al*., 1986; Van Hoesen *et al*., 1991). Since our present neuronal volume fraction estimations are based on semithin sections, and different types of neurons have not been identified, we cannot rule out a selective neuronal loss that may affect either to excitatory neurons or to GABAergic interneurons or to both cell types.

The significantly higher volume fraction occupied by glial somata found in layer II is in line with previous reports showing a larger density of glial cells in human EC from AD cases (Muramori *et al*., 1998). However, no significant differences were found in layer III, suggesting a layer-selective change. The effects of gliosis during AD development are still under debate. It has been proposed that gliosis mediates an inflammatory response to prevent the progression of the disease but that this response may in fact contribute to neurodegeneration (reviewed in Fakhoury, 2018).

### Synaptic changes in AD

#### Synaptic density, proportions and spatial distribution

Our main finding was a significantly lower synaptic density (around 40% lower) in the neuropil of both layer II and layer III of AD individuals compared to controls.

In brain tissue samples from individuals with AD, a lower density of synapses has been described compared to controls in numerous cortical regions, including the DG and certain frontal, parietal, temporal and cingulate cortical areas (Scheff and Price, 1993, 1998, 2003, 2006; Scheff *et al*., 1990, 1996). A decrease in the total number of synapses —accompanied by brain atrophy— has been also reported in CA1, the DG and some cingulate and temporal cortices (Scheff *et al*., 2006, 2007, 2011, 2015; Montero-Crespo *et al*., 2021). However, other studies have not found differences in synaptic density in layers III and V of EC samples from AD cases (Scheff *et al*., 1993). These discrepancies may be due to differences in the methodologies used.. Furthermore, numerous studies have estimated the synaptic density using indirect methods — either using light microscopy to count immunoreactive puncta for synaptic markers (Honer *et al*., 1992; Masliah *et al*., 1990), or employing conventional transmission EM to examine single or few serial ultrathin sections (Scheff *et al*., 1993). Quantification of synaptic density in single ultrathin sections using transmission EM to infer 3D characteristics of synaptic junctions observed in two dimensions may lead to inaccurate synaptic density estimations (see Merchán-Pérez *et al*., 2009). Thus, the present results revealing a lower number of synapses per volume of neuropil, along with the atrophy of the EC compared to controls, suggest a dramatic decrease in the absolute number of synapses in AD cases in these EC layers.

Synaptic loss has been widely reported as the characteristic that best correlates with cognitive deficit in AD patients (reviewed in Colom-Cadena *et al*., 2020). Synaptic loss occurs in the early stages of the disease, affecting firstly the subcortical regions and the EC, and progressing to other cortical regions (Braak and Braak, 1991; Braak and Del Tredici, 2012). Tau protein and β-amyloid peptide in pathological conditions could have toxic effects on synapses leading to synaptic loss and/or dysfunction in AD (Zhou *et al*., 2017; reviewed in Henstridge *et al*., 2016; Rajmohan and Reddy, 2017). This synaptic loss and impairment may cause dysfunction in the cortical circuits contributing to the decline in cognition (reviewed in Colom-Cadena *et al*., 2020).

Previous studies have shown that the percentage of AS and SS varies —80–95% and 20–5%, respectively— in all the cortical layers, cortical areas and species examined so far by TEM (Beaulieu and Colonnier, 1985; Megías et al., 2001; Bourne and Harris, 2011; DeFelipe 2011, 2015). Estimations with samples from human brain using the same method have shown that the AS:SS ratio in layer III of the temporal area 21 is 93:7 (Cano-Astorga et al., 2021), 96:4 in the transentorhinal cortex (Domínguez-Álvaro et al., 2018), and 95:5 in the human CA1 hippocampal field (except in the stratum lacunosum moleculare, in which this ratio was 89:11; Montero-Crespo et al., 2020). Thus, it would appear that the present AS:SS ratio data provide among the highest and lowest proportions previously observed in different brain regions, for AS and SS, respectively. From a functional point of view, the proportion of excitatory and inhibitory synapses is critical, since higher or lower proportions are linked to differences in the excitatory/inhibitory balance of the cortical circuits (for reviews see Froemke, 2015; Zhou and Yu, 2018; Sohal and Rubenstein, 2019). Interestingly, no changes in the proportions of AS and SS were observed; therefore, the lower synaptic density in AD cases might affect AS and SS equally, which is in line with our previous results in the transentorhinal cortex (Domínguez-Álvaro *et al*., 2018). Since most of the synapses are excitatory synapses (∼92% AS), the synaptic decrease may result in a massive loss of AS in particular. As the present results did not reveal differences in the AS:SS ratio between control and AD cases in the neuropil in any layer, it could be interpreted that the proportion of AS and SS on the dendritic arbor of the different types of neurons may be similar. However, it has been shown that there are differences in the number of GABAergic and glutamatergic synaptic inputs in different neuronal types in other cortical regions of a variety of species (e.g., DeFelipe and Fariñas, 1992; Freund and Buzsáki, 1996; DeFelipe, 1997; Somogyi et al., 1998; Schubert et al., 2007; Markram et al., 2015; Tremblay et al., 2016; Hsu et al., 2017). Thus, it would be necessary to examine the synaptic inputs on each specific neuronal type to determine actual differences in the AS:SS ratio in particular cell types, although the final general AS:SS ratio does not vary in the neuropil.

Furthermore, it has been reported that GABA inhibitory interneurons in the brains of AD cases are abnormally reduced, along with the inhibitory neurotransmitters (reviewed in Xu et al., 2020). Other studies have shown that somatostatin neurons, calretinin neurons, and PV neurons are decreased in several brain regions including the EC of AD cases (Solodkin et al., 1996; Mikkonen et al., 1999). Therefore, further studies should be performed to examine the GABAergic innervation of neurons in the EC.

Finally, the present results indicate that the distribution of synapses in the neuropil of both control and AD samples is nearly random, only constrained by the fact that synapses cannot physically overlap in space and so their geometric centers or centroids cannot be too close to their neighbors. As we have previously shown in human brain samples from hippocampal CA1, entorhinal cortex, transentorhinal cortex and temporal cortex, there seems to be no limitation to the position of any synapse except the space already occupied by other synapses (Montero-Crespo et al., 2020; Domínguez-Álvaro et al., 2018, 2021; Cano-Astorga et al., 2021). Previous studies in the plaque-free neuropil of AD cases also showed a random distribution pattern of the synapses in other brain regions (Blazquez-Llorca *et al*., 2013, Domínguez-Álvaro *et al*., 2018; Montero-Crespo *et al*., 2021). It has been reported that during cortical development, synapses are randomly added and/or withdrawn from the population, as described by the synaptic turnover (Rakic et al., 1994). It seems that the only constraint for a synapse to form is that this particular spot is not already occupied by a preexisting synapse (reviewed in Merchán-Pérez et al., 2014). Interestingly, in the AD cases, the loss of synapses per unit volume does not affect their 3D distribution.

#### Shape and size of the synapses

There are very few studies on human brain at the ultrastructural level that provide 3D data on the morphological characteristics of the synaptic junctions to compare with our findings. In the examined EC layers, excitatory AS were larger than inhibitory SS, as previously shown in human transentorhinal cortex (Domínguez-Álvaro *et al*., 2018) and in several strata from the hippocampal CA1 (Montero-Crespo *et al*., 2020). Furthermore, no differences between layers were found regarding the SAS area of AS in AD cases (136,166 nm^2^ in layer II and 122,727 nm^2^ in layer III), which was in contrast with previous results in control EC showing larger synapses in layer III (Domínguez-Álvaro *et al*., 2021). However, the synaptic size in AD samples differs from that of control individuals, with larger and more complex synapses in layer II and a greater proportion of smaller synapses in layer III.

Several EM studies carried out with samples of human brain tissue have reported an increase in the size of synaptic apposition, and this has been suggested as a possible compensatory mechanism for the synaptic loss (reviewed in Scheff and Price, 2006). Models studying the relationship between the size of synaptic junctions and the probability of neurotransmitter release suggest that larger synapses (with a greater number of postsynaptic receptors, mainly AMPA receptors) may produce more powerful, homogeneous and numerous synaptic responses, while smaller synapses (with fewer receptors, mainly NMDA) may produce weaker and more variable responses (Kharazia and Weinberg, 1999; Montes *et al*., 2015). Thus, alterations in the normal size of the synaptic junction may change the proportion of postsynaptic receptors, thereby modifying the synaptic response.

Regarding synaptic shape, we found a higher proportion of horseshoe synapses in layer II from AD cases. In layer III from these AD individuals, higher proportion of AS with macular and fragmented shape were found along with a lower proportion of perforated AS synapses. Several studies have linked an increase in synaptic complexity to an increase in the efficiency of synaptic transmission, via the insertion of new receptors in the postsynaptic membrane (Lüscher *et al*., 2000). Perforated synapses have higher immunoreactivity for glutamate receptors than non-perforated synapses (Ganeshina *et al*., 2004a, *b*). The larger proportion of macular AS in layer III (around 83% in AD cases, versus 76% in control cases) and the corresponding lower number of perforated synapses (about half the number found in control cases) supports the evidence for a larger number of simple synapses in this layer in AD cases, which is in turn supported by the finding of a higher proportion of smaller synapses described above. However, the higher proportion of fragmented AS in layer III from AD cases may indicate the above-mentioned compensatory response to the synaptic loss. Conversely, small macular synapses from layer III may be more susceptible to damage resulting in a reduction in their numbers.That is, synaptic morphology may be selectively altered by layer, with a non-homogenous pattern of changes, which would differentially affect the circuits in which these layers are involved.

Therefore, synaptic changes in layer II and III might be part of a remodeling process of the remaining synapses, which overcomes the overall synaptic loss. If the remodeling process involves an increase in the proportion of synapses with more complex morphology, it could be hypothesized that the morphological changes in AD would trigger compensatory mechanisms during the progression of the disease. However, this type of compensatory mechanism may fail or be deficient, leading to a decline in the synaptic functionality accompanying the progression of the disease.

#### Postsynaptic targets

Despite the lower synaptic density in AD cases, no changes in the distribution and types of postsynaptic targets were found. In the AD samples, the most frequent combinations were axospinous AS (on dendritic spine heads) and axodendritic AS (established on dendritic shafts). In addition, the majority of the synapses that a pyramidal cell receives are on spines and they represent the vast majority of AS synapses (DeFelipe and Fariñas, 1992). In the human transentorhinal cortex, 75% of the synapses were AS on spines, which is similar to the case of the AS found in the layer III of the temporal cortex (75%) using the same analysis method (Cano-Astorga et al., 2021). This percentage was much higher in the CA1 hippocampal field, as high as 94% in the superficial part of the CA1 stratum pyramidale (Montero-Crespo et al., 2020). In both layers II and III of the entorhinal cortex, the percentage of AS on spines was 57%, and 43% in the case of AS on dendritic shafts. This is a higher proportion of AS on shafts than in the other human cortical regions previously analyzed using the same techniques. In addition, the comparison between control and Alzheimer’s disease samples did not show a clear reduction in the proportion of AS on spines. Only a slightly lower proportion of AS established on spines was found in layer II from Alzheimer’s disease samples. Therefore, the lower synaptic density does not seem to be specifically related to a loss of spines, but rather a general loss of synapses might occur regardless of their postsynaptic targets. It may be that the axospinous synapses in EC from AD cases are functionally altered even though no changes were found in their proportions.

#### Implications of the synaptic changes

EC provides excitatory inputs to the hippocampus from layer II neurons targeting the dentate gyrus (trisynaptic pathway) and from layer III neurons monosynaptically targeting CA1 (reviewed in Insausti and Amaral, 2012; Marks et al., 2020). Neuronal projections from both layers give rise to the perforant pathway, which has generally been reported to be weakened in AD (e.g., Hyman *et al*., 1986).

Moreover, in layer II there are two subpopulations of excitatory neurons: spiny stellate neurons (which project to DG, CA3 and CA2) and modified pyramidal neurons (which project to stratum lacunosum in CA1). Projections from these neurons, constituting the trisynaptic pathway, seem to be related to contextual memories (reviewed in Mark et al., 2020). It has been proposed that this pathway is more susceptible to premature degeneration than the monosynaptic pathway (Van Hoesen et al., 2006; Llorens-Martín et al., 2014).

In addition, pyramidal neurons from layer III directly project to pyramidal neuron dendrites located in the *stratum moleculare* of CA1, and it has been proposed that these monosynaptic excitatory inputs drive temporal andradetion learning (reviewed in Mark et al., 2020).

The present study shows a significant lower number of synapses per volume, as well as morphological synaptic alterations in the EC from AD cases in layers II and III. Since the AD cases examined in the present study correspond to advanced stages of the disease, we do not know *when* the synaptic alterations occurred. Considering that there is neuronal loss, the lower number of synapses may be explained —at least in part— by the loss of local axons of these neurons, at least at the earlier stages of the disease. However, the lower number of synapses may also be explained by a loss of projection neurons that die in distant cortical regions that are also affected by the disease.

#### Interindividual variability

The present data cannot be generalized to the whole population of patients with AD, which shows clear variability between individuals (Fig. 7; Table 2). Therefore, our study can be considered as a further step to tackle the issue of the synaptic alterations in Alzheimer’s disease cases, but it would be necessary to validate our results, both in a larger number of individuals and in additional brain regions (which we have already done in the hippocampal CA1 field and in the trans-entorhinal cortex). Technical effects could be ruled out given that the postmortem delays were all similar and the procedures used were the same. Moreover, all of the AD cases examined were women aged between 80 and 86 years old; therefore, variability due to sex and aging could also be ruled out.

Considering the above-mentioned toxic effect of tau protein and β-amyloid peptide on the synapses, a relationship between the degree of disease progression and the synaptic loss might be expected: the greater the degree of the pathology, the greater the synaptic loss. Subject VK16, who presented a high score for neuropathology and disease progression (Braak/CERAD: VI/C), also had the lowest synaptic density in both layers (0.12 synapse/µm^3^; Fig. 7; Extended Table 2-1). However, subject VK22, who also had a high score for neuropathology and disease progression (V/C), had a synaptic density of more than double (0.29 and 0.27 synapse/µm^3^, for layers II and III, respectively) and both cases, VK16 and VK22, suffered dementia. However, subject IF1 did not present evidence of dementia despite the fact that this subject had a high level of pathology (IV/B) and low synaptic density, which was similar to VK22 (0.21 and 0.27 synapse/µm^3^, for layers II and III, respectively). However, the cortical thickness of case IF1 was 2.3 mm, whereas —in VK16 and VK22— the cortical layers were thinner (1.19 and 1.51 mm, respectively; Extended Table 1-1). As pointed out by Ferrer (2012), it should be kept in mind that AD —at least and limited to the entorhinal and transentorhinal cortices (stages I–II)— affects about 80% of individuals over 65 years, but dementia only occurs in a small percentage of individuals in this age bracket (the prevalence of dementia in AD increases to 25% in 80-year-old individuals). Thus, it is possible that this particular case (IF1) may represent a pre-dementia stage of AD (prodromal AD).

Control cases were on average younger (51 years old) than Alzheimer’s disease cases (85 years old) and it remains uncertain whether some of the synaptic alterations observed in the present work were due to normal aging or due to the pathology per se. Although no sex-based differences have been reported with aging in the human cerebral cortex (Scheff *et al*., 2001), previous studies on synaptic density in the human *temporal* cortex have shown differences depending on the sex (Alonso-Nanclares *et al*., 2008). Nevertheless, no similar EM studies have been performed in the EC in individuals with different ages and sexes. Thus, the lower synaptic density found in AD cases might also be the results of sex or age effects. In aged rhesus monkey, a lower number of synapses has been reported in prefrontal cortex related to a cognitive decline; however, studies in rats and monkeys have shown no evidence of synaptic loss with age in mesial temporal lobe structures (reviewed in Morrison and Baxter, 2012). Further studies using human brain tissue from older control cases would be necessary to determine whether or not these differences in the synaptic density are due to these factors, as opposed to the AD *per se*. Nevertheless, it is important to point out that only the neuropil without plaques was examined in the present study, whereas it is well established that the neuropil which is adjacent to —and is surrounding— the plaques shows synaptic alterations (Blazquez-Llorca *et al*., 2013). Thus, it is possible that the neuropil lacking plaques in AD cases displays “normal” characteristics, and differences observed in our study comparing to the control group might be due to factors related to sex and age.

## Conclusions

In the present study, we found, in addition to synaptic loss, changes in the morphology of the synapses in AD compared to control cases. These structural changes may contribute to the anatomical basis for the impairment of cognitive functions in AD. Despite the large number of synapses analyzed (5000 synapses), and the proven robustness of the methodology used in this study, it should be kept in mind that (i) the data comes from the analysis of 4 control cases and 4 AD cases and (ii) we cannot rule out sex-and age-related factors that may influence the differences observed between the two groups. In order to address these remaining issues, it would be necessary to perform further studies using FIB/SEM or other similar techniques to reconstruct thousands of synapses in aged control cases of both sexes.

## Declarations

### Availability of data and materials

Most data generated or analyzed during this study are included in the main text and the tables. The control datasets used and analyzed are available on the EBRAINS Knowledge Graph (DOI: 10.25493/3GMJ-FEZ). Datasets used during the current study available from the corresponding author on reasonable request.

## Authors’ contributions

M D-A, Conceptualization, Data curation, Software, Formal analysis, Methodology, Writing -review and editing; M M-C, Data curation, Software, Methodology; L B-L, Conceptualization, Supervision, Validation, Methodology, Writing - review and editing; S P-A, Data analysis and curation, Writing - review and editing; N C-A, Data curation, Software, Writing - review and editing; J DeF, Conceptualization, Resources, Supervision, Funding acquisition, Validation, Project administration, Writing – review and editing; L A-N, Conceptualization, Data curation, Formal analysis, Supervision, Validation, Investigation, Methodology, Writing - original draft, Writing - review and editing.

## Acknowledgments

We would like to thank Carmen Álvarez and Lorena Valdés for their technical assistance, and Nick Guthrie for his excellent text editing.

## Extended data legends

**Movie 1. EspINA software user interface.** FIB/SEM sections are viewed through the *xyz*-planes. Asymmetric (green) synapses are shown — established on dendritic spines originating from a dendritic shaft (purple), shown as D529 in Figure 5.

**Movie 2. EspINA software 3D viewer.** A stack of images is fully represented in the three orthogonal planes (x, y and z). The video shows the 3D reconstruction of a single dendritic segment (purple; D529 shown in Figure 5) and the 3D reconstruction of all asymmetric (green) and symmetric (red) synapses established in this 3D reconstructed fragment.

**Extended Figure 9-1.**
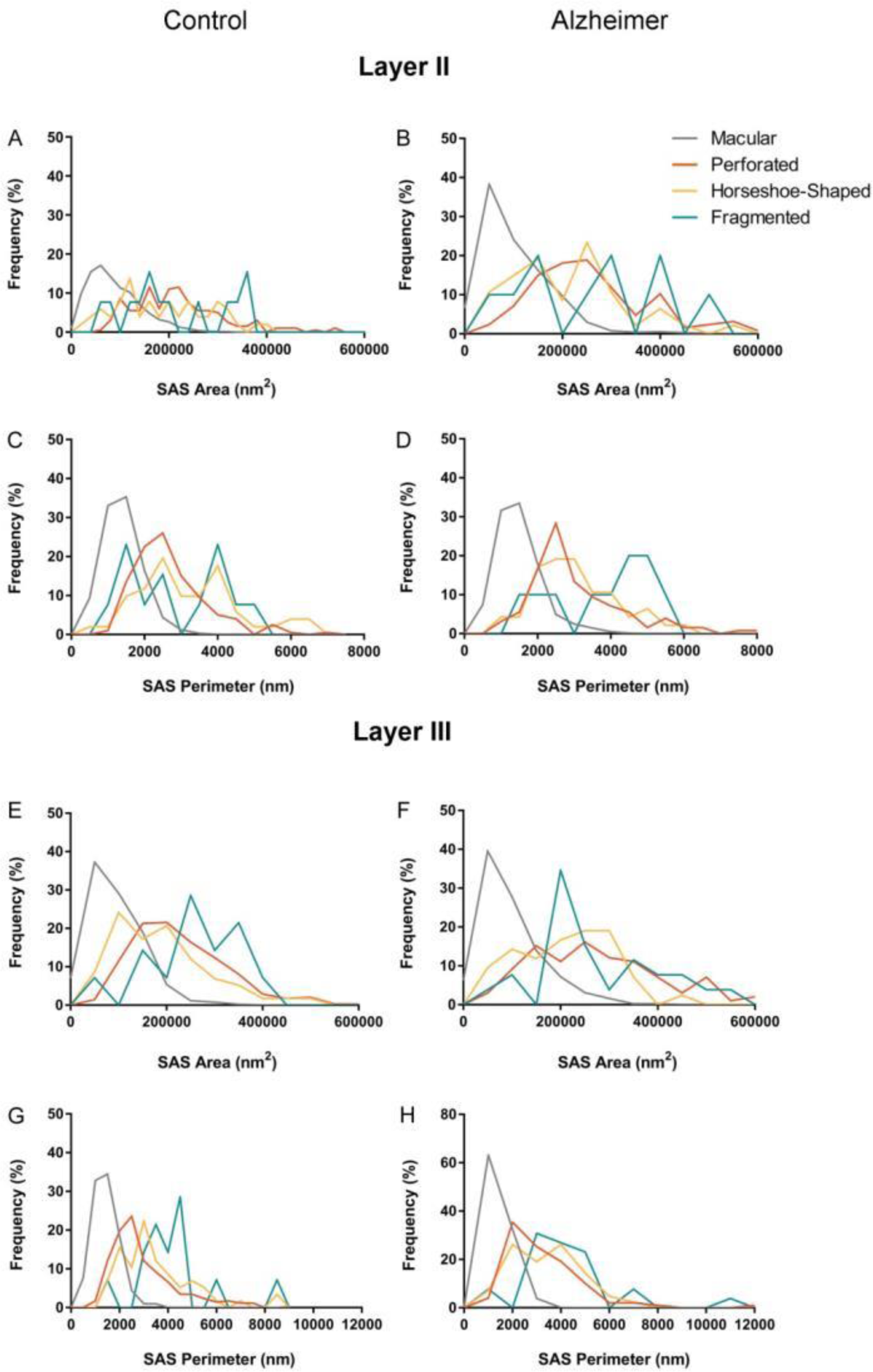
Frequency histograms of the SAS area (A, B, E, F) and perimeter (C, D, G, H) of macular, perforated, horseshoe-shaped and fragmented AS from control cases (A, C, E, G) and from AD cases (B, D, F, H), in layers II and III of the EC. The SAS area and perimeter of macular synapses were significantly smaller than in perforated, horseshoe-shaped and fragmented synapses (KW, p<0.0001). No differences were found between control and AD cases (KS, p>0.0001).

**Extended Figure 10-1.**
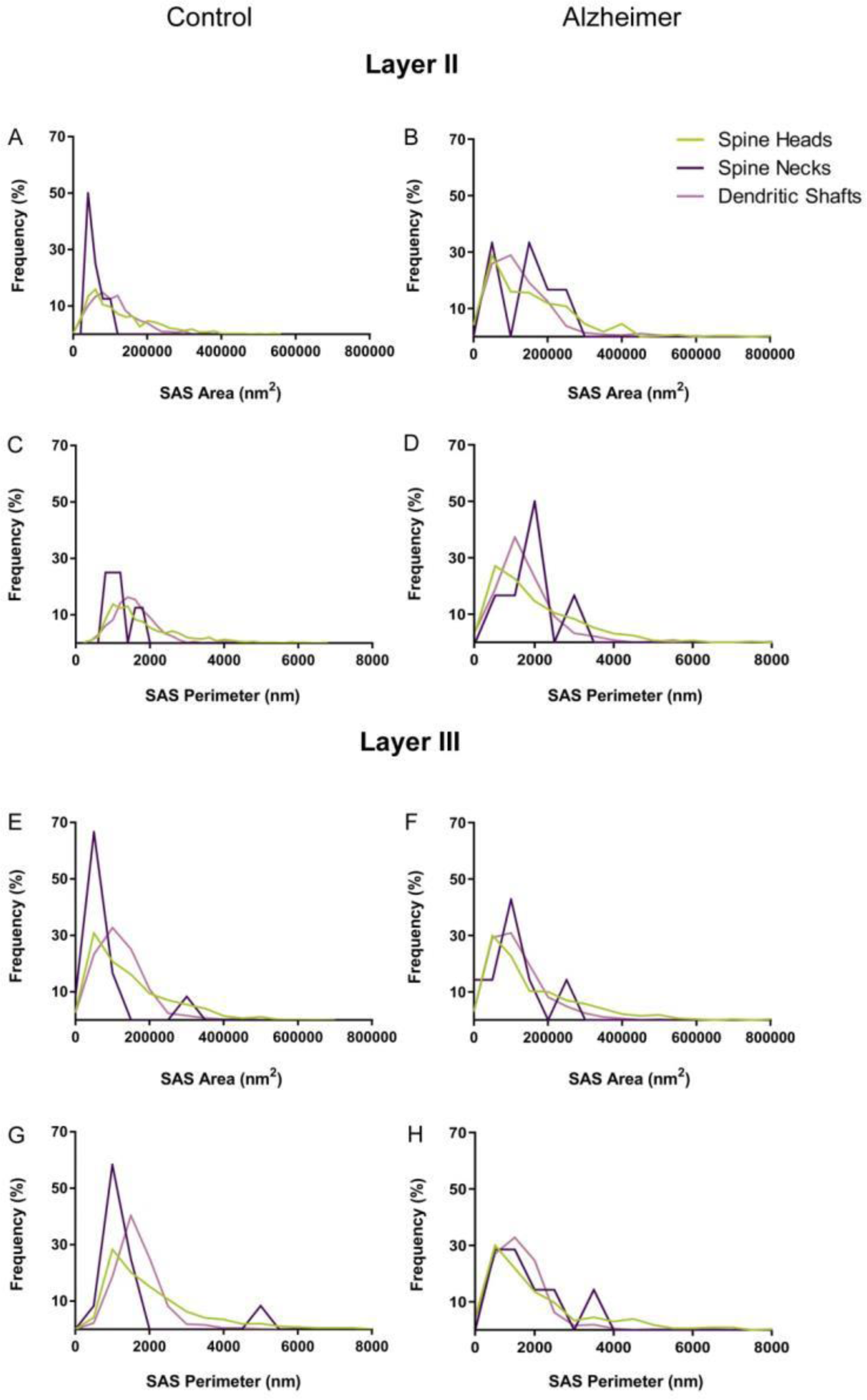
Frequency histograms of the SAS area (A, B, E, F) and perimeter (C, D, G, H) of AS targeting spine heads, spine necks and dendritic shafts from control cases (A, C, E, G) and from AD cases (B, D, F, H), in both layers II and III of the EC. No differences were found between groups (KS, p>0.0001).

**Extended Table 1-1.** EC cortical thickness in control and Alzheimer cases. All data are corrected for shrinkage factor.

| **Group** | **Case** | **EC thickness** |
| --- | --- | --- |
|  |  | **(mm; mean±**<br>**SD)** |
| **Control** | **AB1** | 2.12±0.06 |
|  | **AB3** | 2.08±0.01 |
|  | **IF10** | 1.75±0.05 |
|  | **M16** | 2.04±0.02 |
| **Alzheimer** | **IF1** | 2.30±0.03 |
|  | **VK11** | 1.40±0.01 |
|  | **VK16** | 1.19±0.01 |
|  | **VK22** | 1.51±0.02 |

**Extended Table 1-2.** Volume fraction occupied by cortical elements in layers II and III of the EC. Data are given as mean±SD. EC: entorhinal cortex; SD: standard deviation; V_neu_: volume fraction occupied by neurons; V_g_: volume fraction occupied by glia; V_bv_: volume fraction occupied by blood vessels; V_n_: volume fraction occupied by neuropil. Data from individual cases are shown in Extended Table1-3.

| **EC**<br>**layer** | **Group** | **V<sub>neu</sub> (%)** | **V<sub>g</sub> (%)** | **V<sub>bv</sub> (%)** | **V<sub>n</sub> (%)** |
| --- | --- | --- | --- | --- | --- |
| **II** | **Control** | 4.49±2.91 | 2.64±1.05 | 3.80±2.31 | 89.07±3.67 |
|  | **Alzheimer** | 2.61±1.41 | 4.59±0.90 | 2.34±1.59 | 90.47±2.03 |
| **II** | **Control** | 7.53±1.78 | 2.52±0.42 | 3.80±0.74 | 86.15±1.23 |
|  | **Alzheimer** | 2.88±1.48 | 4.55±1.85 | 3.98±2.73 | 88.59±3.94 |

**Extended Table 1-3.**
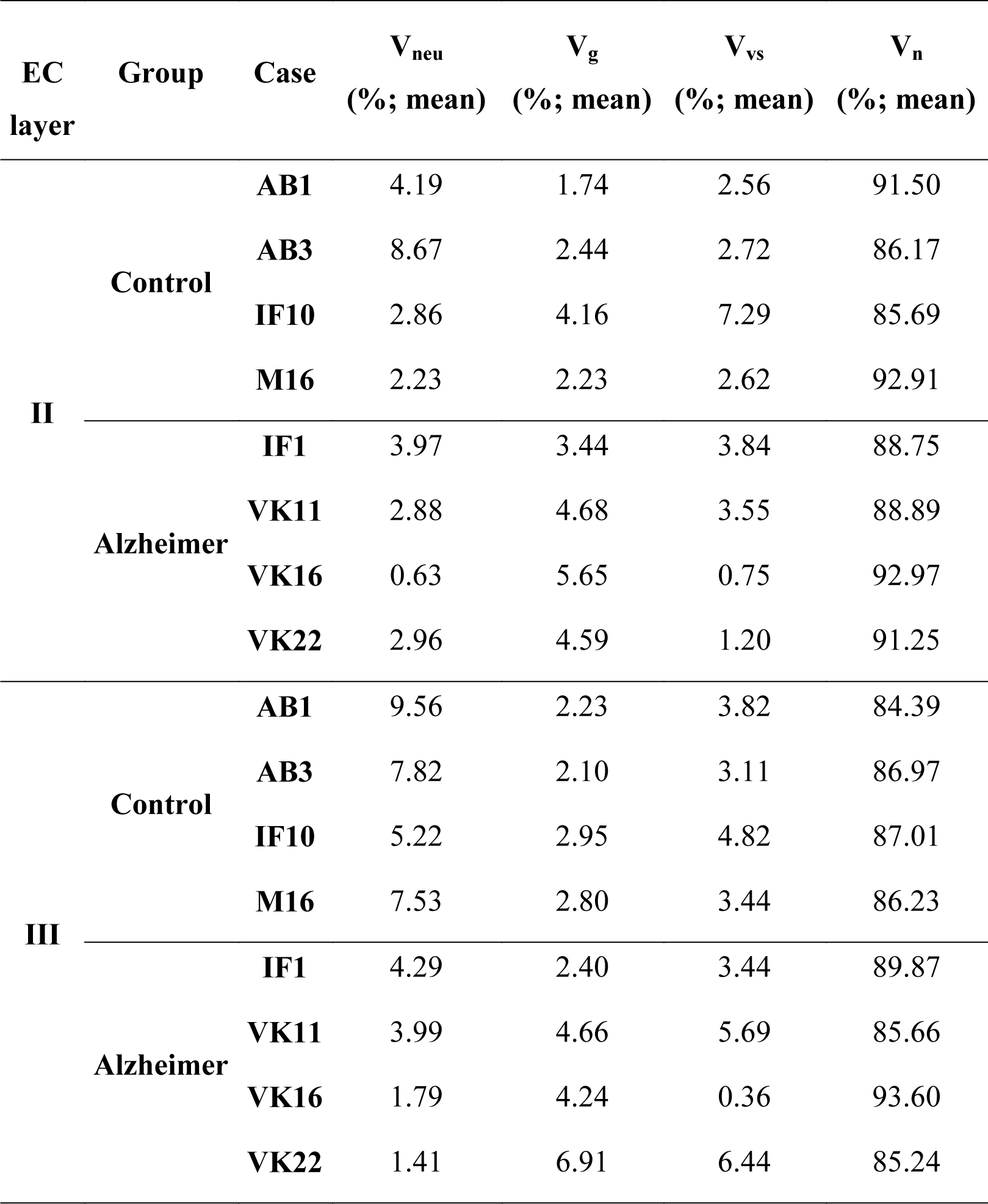
Light microscopy data on volume fraction occupied by cortical elements in layers II and III of the EC for individual cases. All volume data are corrected for shrinkage. EC: entorhinal cortex; V_neu_: volume fraction occupied by neurons; V_g_: volume fraction occupied by glia; V_bv_: volume fraction occupied by blood vessels; V_n_: volume fraction occupied by neuropil.

**Extended Table 2-1.**
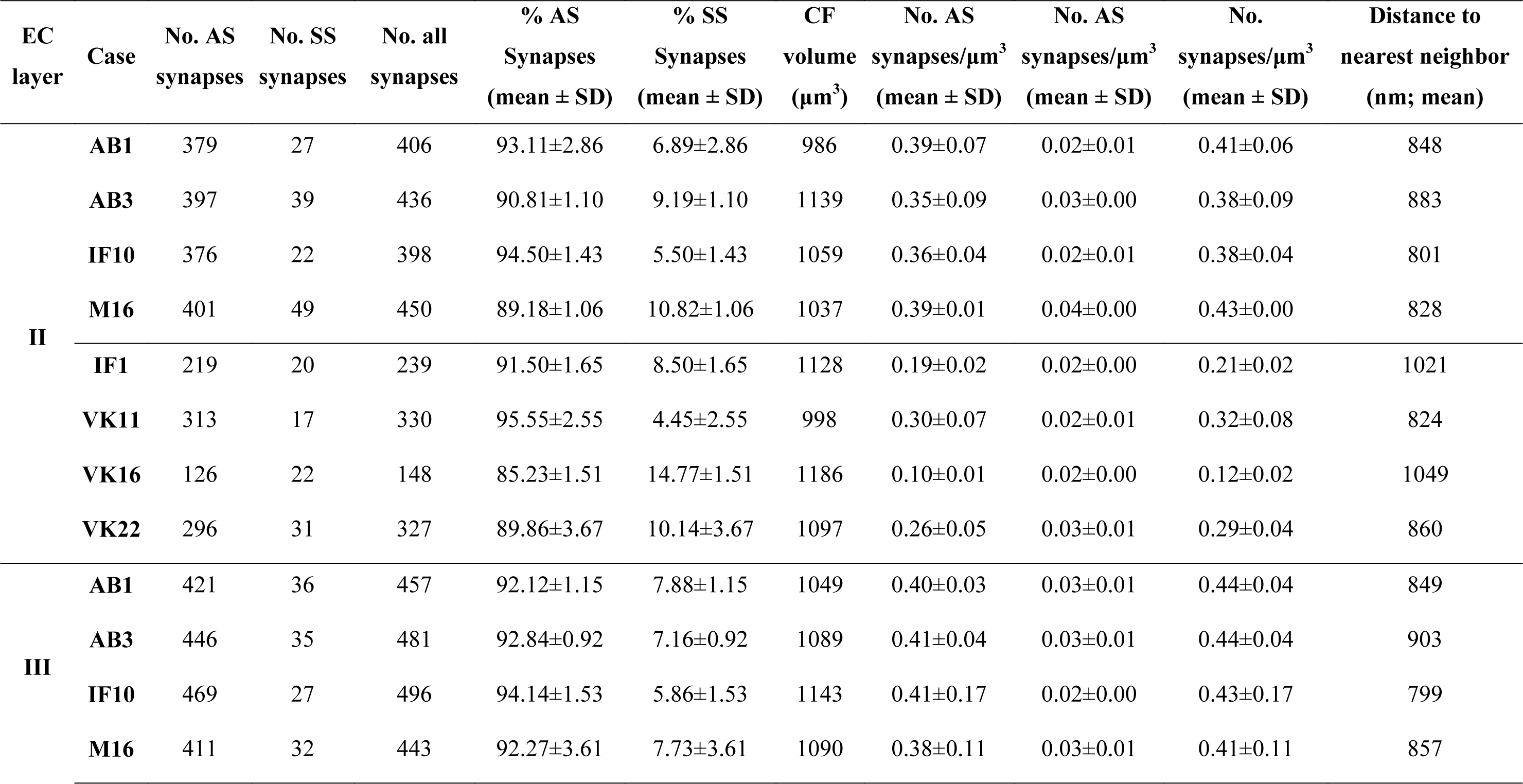
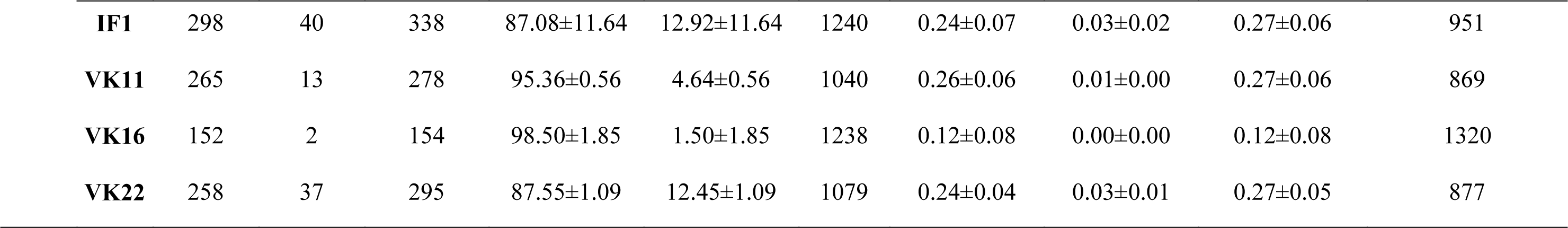
Data from the ultrastructural analysis of neuropil from layers II and III of the EC for individual cases. All volume and distance data are corrected for shrinkage. All: includes AS+SS synapses; AS: asymmetric synapses; EC: entorhinal cortex; No.: number; SD: standard deviation; SS: symmetric synapses.

**Extended Table 3-1.** Area (nm^2^) and perimeter (nm) of the SAS in layers II and III of the EC for individual cases. All data are corrected for shrinkage factor. AS: asymmetric synapses; EC: entorhinal cortex; sem: standard error of the mean; SAS: synaptic apposition surface; SS: symmetric synapses.

| Case | Type of synapse | Layer II |  | Layer III |  |
| --- | --- | --- | --- | --- | --- |
|  |  | Area of SAS | Perimeter of SAS | Area of SAS | Perimeter of SAS |
| **AB1** | AS | 101,766±3757 | 1551±40 | 116,843±3,974 | 1673±43 |
|  | SS | 58,790±6865 | 1252±117 | 60,978±5,015 | 1220±79 |
| **AB3** | AS | 109,320±3694 | 1605±38 | 132,252±4,051 | 1859±47 |
|  | SS | 71,082±13,154 | 1593±226 | 67,083±5,083 | 1444±89 |
| **IF10** | AS | 113,399±4343 | 1770±51 | 121,736±4,345 | 1991±60 |
|  | SS | 53,681±4642 | 1331±112 | 65,338±9,444 | 1578±210 |
| **M16** | AS | 116,757±4139 | 1598±40 | 125,900±4,790 | 1779±55 |
|  | SS | 80,435±6621 | 1440±73 | 76,379±10,422 | 1354±129 |
| **IF1** | AS | 163,933±7143 | 1909±54 | 146,264±5,934 | 1975±72 |
|  | SS | 77,503±12,221 | 1422±141 | 99,073±8,446 | 2149±170 |
| **VK11** | AS | 120,779±5614 | 1858±66 | 134,348±7,330 | 1858±83 |
|  | SS | 53,824±6035 | 1236±80 | 46,814±13,543 | 1161±254 |
| **VK16** | AS | 142,393±15,315 | 1841±131 | 106,568±7,258 | 1511±53 |
|  | SS | 46,667±5245 | 1070±77 | 46,515±7,167 | 1103±20 |
| **VK22** | AS | 117,558±5198 | 1668±50 | 103,726±5,248 | 1536±52 |
|  | SS | 58,932±6912 | 1227±87 | 36,680±2,946 | 959±56 |

**Extended Table 4-1.** Proportion of macular, perforated, horseshoe-shaped and fragmented synapses in layer II of the EC for individual cases. Data are given as percentages with the absolute number of synapses studied in parentheses. AS: asymmetric synapses; EC: entorhinal cortex; SS: symmetric synapses.

| **Case** | **Type of synapse** | **Macular synapses** | **Perforated synapses** | **Horseshoe synapses** | **Fragmented synapses** | **Total synapses** |
| --- | --- | --- | --- | --- | --- | --- |
| **AB1** | AS | 84.7% (320) | 9.8% (37) | 4.5% (17) | 1.0% (4) | 100% (378) |
|  | SS | 74.1% (20) | 7.4% (2) | 18.5% (5) | 0% (0) | 100% (27) |
| **AB3** | AS | 82.6% (327) | 13.6% (54) | 2.5% (10) | 1.3% (5) | 100% (396) |
|  | SS | 66.6% (26) | 15.4% (6) | 15.4% (6) | 2.6% (1) | 100% (39) |
| **IF10** | AS | 81.6% (307) | 14.1% (53) | 4.0% (15) | 0.3% (1) | 100% (376) |
|  | SS | 72.7% (16) | 9.1% (2) | 18.2% (4) | 0% (0) | 100% (22) |
| **M16** | AS | 83.0% (331) | 14.0% (56) | 2.3% (9) | 0.7% (3) | 100% (399) |
|  | SS | 77.6% (38) | 10.2% (5) | 12.2% (6) | 0% (0) | 100% (49) |
| **IF1** | AS | 86.8% (190) | 9.6% (21) | 3.2% (7) | 0.4% (1) | 100% (219) |
|  | SS | 70.0% (14) | 15.0% (3) | 15.0% (3) | 0% (0) | 100% (20) |
| **VK11** | AS | 77.3% (242) | 14.7% (46) | 6.1% (19) | 1.9% (6) | 100% (313) |
|  | SS | 58.8% (10) | 23.5% (4) | 11.8% (2) | 5.9% (1) | 100% (17) |
| **VK16** | AS | 79.4% (100) | 19.0% (24) | 1.6% (2) | 0% (0) | 100% (126) |
|  | SS | 77.3% (17) | 0% (0) | 18.2% (4) | 4.5% (1) | 100% (22) |
| **VK22** | AS | 80.1% (237) | 12.5% (37) | 6.4% (19) | 1.0% (3) | 100% (296) |
|  | SS | 80.6% (25) | 12.9% (4) | 6.5% (2) | 0% (0) | 100% (31) |

**Extended Table 4-2.**
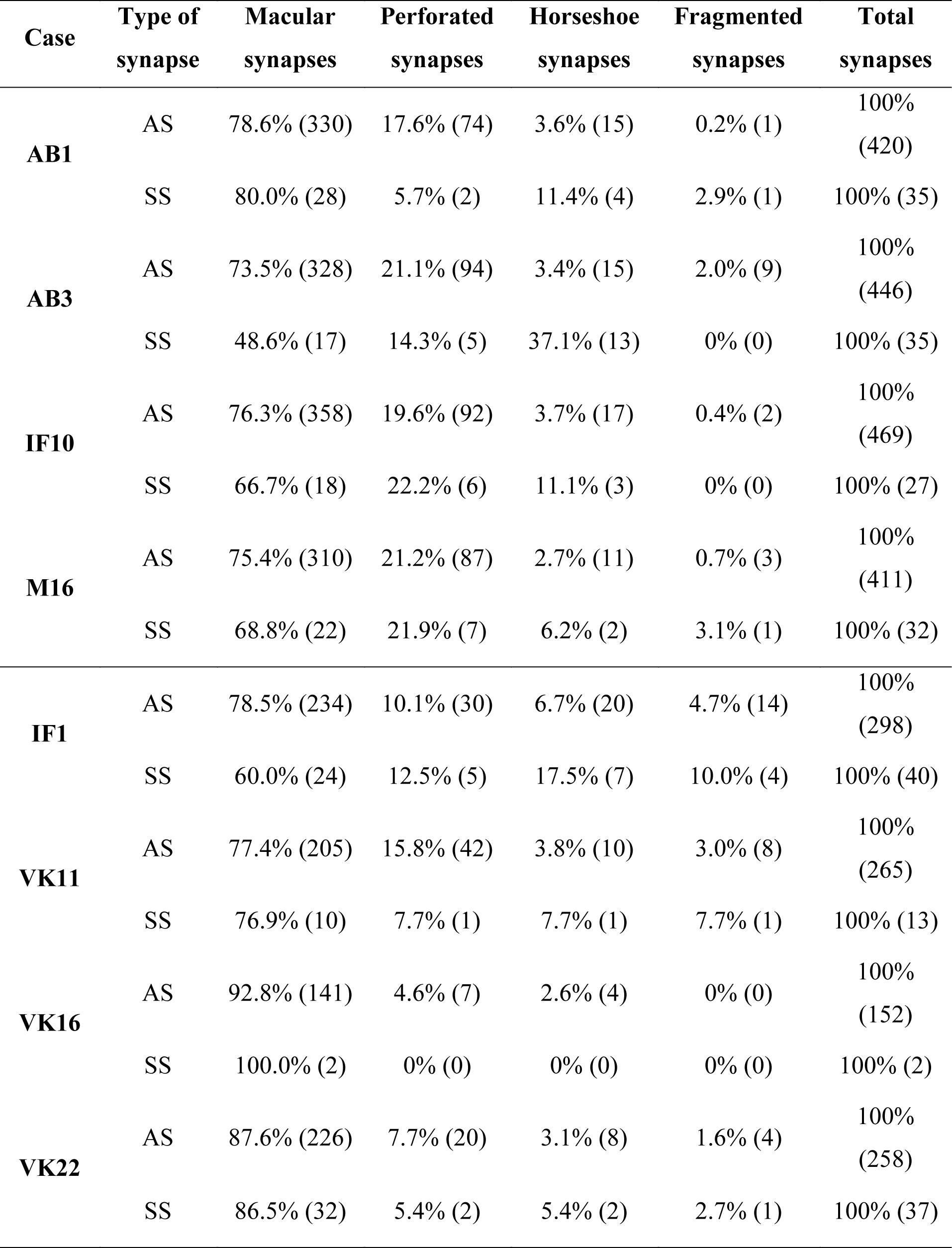
Proportion of macular, perforated, horseshoe-shaped and fragmented synapses in layer III of the EC for individual cases. Data are given as percentages with the absolute number of synapses studied in parentheses. AS: asymmetric synapses; EC: entorhinal cortex; SS: symmetric synapses.

**Extended Table 4-3.** Area (nm^2^) and perimeter (nm) of the macular, perforated, horseshoe-shaped and fragmented SAS in layer II of the EC. All data are corrected for shrinkage. AS: asymmetric synapses; EC: entorhinal cortex; sem: standard error of the mean; SAS: synaptic apposition surface; SS: symmetric synapses.

| **Group** | **Shape** | **Type of synapse** | **SAS Area (nm<sup>2</sup>; mean± sem)** | **SAS Perimeter (nm; mean± sem)** |
| --- | --- | --- | --- | --- |
| **Control** | **Macular** | AS | 89,406±1999 | 1389±21 |
|  |  | SS | 53,716±7280 | 1126±32 |
|  | **Perforated** | AS | 218,803±7373 | 2698±121 |
|  |  | SS | 104,339±22,261 | 1730±413 |
|  | **Horseshoe** | AS | 191,238±14,403 | 3242±273 |
|  |  | SS | 84,034±6931 | 2046±47 |
|  | **Fragmented** | AS | 199,121±19,630 | 2818±269 |
|  |  | SS | 152,108±0 | 2500±0 |
| **Alzheimer** | **Macular** | AS | 101,410±13,147 | 1474±87 |
|  |  | SS | 50,519±4291 | 1101±59 |
|  | **Perforated** | AS | 290,075±33,353 | 3146±261 |
|  |  | SS | 87,027±14,137 | 1584±95 |
|  | **Horseshoe** | AS | 230,872±27,294 | 3296±196 |
|  |  | SS | 95,635±24,799 | 1821±223 |
|  | **Fragmented** | AS | 329,742±94,617 | 3993±782 |
|  |  | SS | 54,638±11,867 | 1323±150 |

**Extended Table 4-4.** Area (nm^2^) and perimeter (nm) of the macular, perforated, horseshoe-shaped and fragmented SAS in layer III of the EC. All data are corrected for shrinkage factor. AS: asymmetric synapses; EC: entorhinal cortex; sem: standard error of the mean; SAS: synaptic apposition surface; SS: symmetric synapses.

| **Group** | **Shape** | **Type of synapse** | **SAS Area (nm<sup>2</sup>; mean± sem)** | **SAS Perimeter (nm; mean± sem)** |
| --- | --- | --- | --- | --- |
| **Control** | **Macular** | AS | 92,737±3846 | 1430±32.73 |
|  |  | SS | 65,303±14,798 | 1110±27.5 |
|  | **Perforated** | AS | 228,057±6993 | 2964±173.7 |
|  |  | SS | 86,820±5148 | 1695±86.54 |
|  | **Horseshoe** | AS | 18,8362±9007 | 3447±340.4 |
|  |  | SS | 157,439±53,309 | 2343±588.2 |
|  | **Fragmented** | AS | 303,754±35,891 | 4254±505.9 |
|  |  | SS | 62,518±5339 | 2583±1624 |
| **Alzheimer** | **Macular** | AS | 96,162±6492 | 1412±47.63 |
|  |  | SS | 47,435±9453 | 1120±156.6 |
|  | **Perforated** | AS | 296,156±24,116 | 3192±150.9 |
|  |  | SS | 99,845±47,349 | 2723±868.4 |
|  | **Horseshoe** | AS | 192,661±19,980 | 3117±283.5 |
|  |  | SS | 127,306±39,972 | 2289±479.0 |
|  | **Fragmented** | AS | 250,707±40,241 | 3577±781.0 |
|  |  | SS | 70,021±43,790 | 1758±908.2 |

**Extended Table 5-1.** Distribution of AS and SS on spines and dendritic shafts in layer II of the EC for individual cases. Synapses on spines have been sub-divided into those that are established on spine heads and those that are established on spine necks. Synapses on spine have been recorded separately. Moreover, we differentiated between aspiny and spiny dendritic shafts. Data are expressed as percentages with the absolute number of synapses studied given in parentheses. AD: Alzheimer’s disease; AS: asymmetric synapses; EC: entorhinal cortex; SS: symmetric synapses.

| **Case** | **Type of synapse** | **Synapses on spine heads** | **Synapses on nf, spine heads** | **Synapses on spine necks** | **Synapses on aspiny dendritic shafts** | **Synapses on spiny dendritic shafts** | **Total synapses** |
| --- | --- | --- | --- | --- | --- | --- | --- |
| AB1 | AS | 35.3% (121) | 20.7% (71) | 0.6% (2) | 26.5% (91) | 16.9% (58) | 100% (343) |
|  | SS | 4.8% (1) | 4.8% (1) | 0.0% (0) | 33.3% (7) | 57.1% (12) | 100% (21) |
| AB3 | AS | 31.6% (116) | 17.9% (66) | 0.6% (2) | 24.8 (91) | 25.1% (92) | 100% (367) |
|  | SS | 10.8% (4) | 5.4% (2) | 0.0% (0) | 27.0% (10) | 56.8% (21) | 100% (37) |
| IF10 | AS | 36.7% (122) | 22.6% (75) | 0.3% (1) | 17.2% (57) | 23.2% (77) | 100% (332) |
|  | SS | 9.1% (2) | 9.1% (2) | 0.0% (0) | 18.2% (4) | 63.6% (14) | 100% (22) |
| M16 | AS | 53.7% (208) | 15.0% (58) | 0.8% (3) | 11.6% (45) | 18.9% (73) | 100% (387) |
|  | SS | 10.4 (5) | 2.1% (1) | 2.1% (1) | 16.7% (8) | 68.7% (33) | 100% (48) |
| IF1 | AS | 34.4% (71) | 15.9% (33) | 0.5% (1) | 21.7% (45) | 27.5% (57) | 100%( 207) |
|  | SS | 20.0% (4) | 5.0% (1) | 0.0% (0) | 35.0% (7) | 40.0% (8) | 100% (20) |
| VK11 | AS | 35.5% (89) | 23.5% (59) | 0.8% (2) | 20.7% (52) | 19.5% (49) | 100% (251) |
|  | SS | 13.3% (2) | 0.0% (0) | 0.0% (0) | 40.0% (6) | 46.7% (7) | 100% (15) |
| VK16 | AS | 27.1% (26) | 21.9% (21) | 3.1% (3) | 32.3% (31) | 15.6% (15) | 100% (96) |
|  | SS | 5.3% (1) | 0.0% (0) | 0.0% (0) | 57.9% (11) | 36.8% (7) | 100% (19) |
| VK22 | AS | 38.7% (98) | 12.3% (31) | 0.4% (1) | 29.6% (75) | 19.0% (48) | 100% (253) |
|  | SS | 7.1% (2) | 0.0% (0) | 0.0% (0) | 57.2% (16) | 35.7% (10) | 100% (28) |

**Extended Table 5-2.**
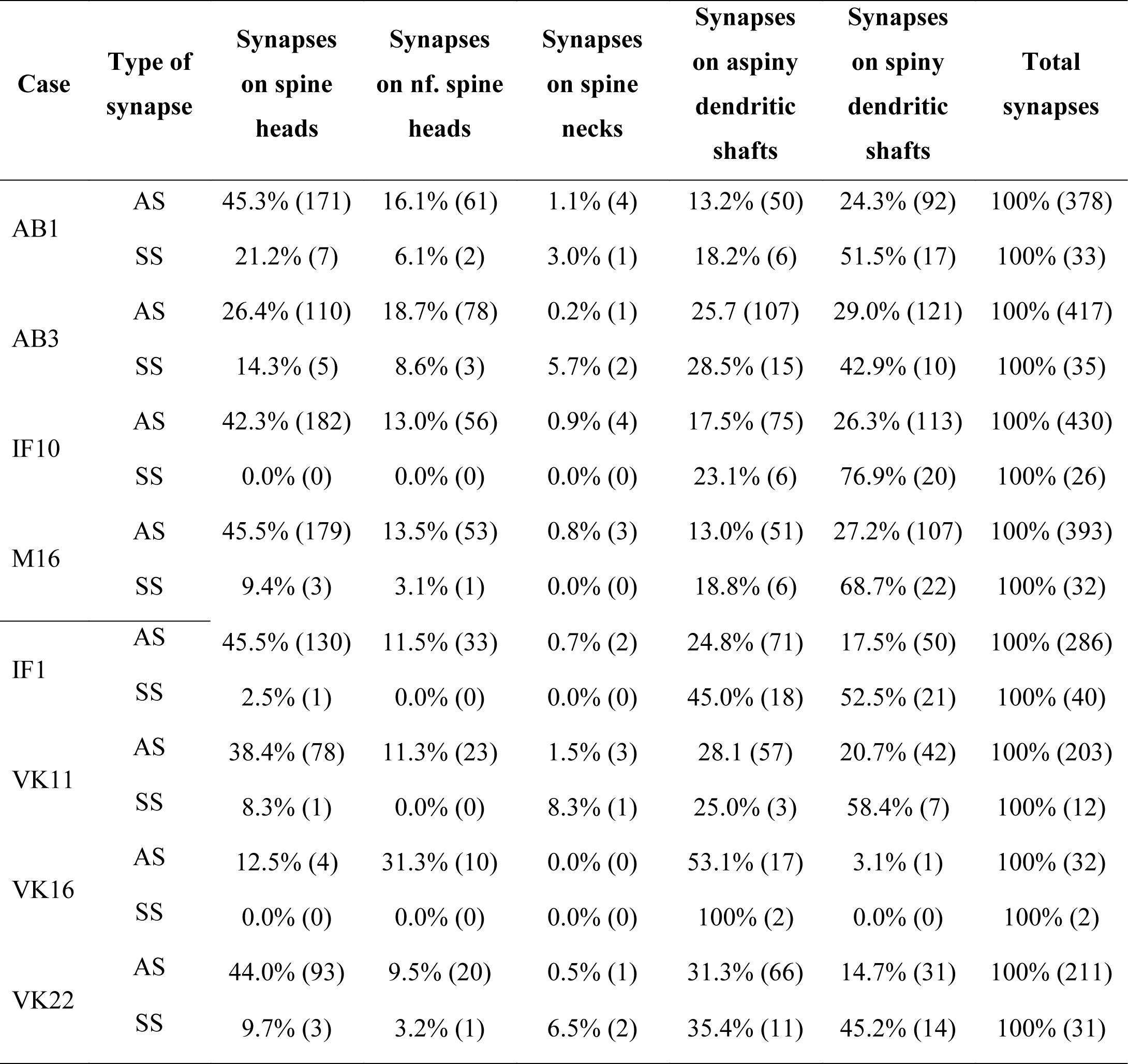
Distribution of AS and SS on spines and dendritic shafts in layer III of the EC for individual cases. Synapses on spines have been sub-divided into those that are established on spine heads and those that are established on spine necks. Synapses on spines have been recorded separately. Moreover, we differentiated between aspiny and spiny dendritic shafts. Data are expressed as percentages with the absolute number of synapses studied given in parentheses. AD: Alzheimer’s disease; AS: asymmetric synapses; EC: entorhinal cortex; SS: symmetric synapses.

